# Identification of potent anti-fibrinolytic compounds against plasminogen and tissue-type plasminogen activator using computational approaches

**DOI:** 10.1101/2022.10.13.512028

**Authors:** Suparna Banerjee, Yeshwanth M, Dhamodharan Prabhu, Kanagaraj Sekar, Prosenjit Sen

**Affiliations:** School of Biological Sciences, Indian Association for the Cultivation of Science, Jadavpur, Kolkata 700032, West Bengal, India; Department of Computational and Data Sciences, Indian Institute of Science, Bangalore 560012, Karnataka, India; Research and Development Wing, Sree Balaji Medical College and Hospital-BIHER, Chennai 600044, Tamil Nadu, India

**Keywords:** plasmin, structure-based virtual screening, molecular docking, molecular dynamics simulation, principal component analysis (PCA), Molecular Mechanics Poisson-Boltzmann Surface Area (MMPBSA)

## Abstract

The zymogen protease Plasminogen (Plg) and its active form plasmin (Plm) carry out important functions in the blood clot disintegration (breakdown of fibrin fibres) process. Inhibition of plasmin effectively reduces fibrinolysis to circumvent heavy bleeding. Currently, available Plm inhibitor tranexamic acid (TXA) that is used to treat severe hemorrhages is associated with an increased incidence of seizures which in turn were traced to gamma-aminobutyric acid antagonistic activity (GABAa) in addition to having multiple side effects. Fibrinolysis can be suppressed by targeting the three important protein domains: kringle-1 and serine protease domain of plasminogen and kringle-2 domain of tissue plasminogen activator. In the present study, combined approaches of structure-based virtual screening and molecular docking using Schrödinger Glide, AutoDock Vina, and ParDock/BAPPL+ were employed to identify potential hits from the ZINC database. Thereafter, the drug-likeness properties of the top three leads for each protein target were evaluated using Discovery Studio. Subsequently, a molecular dynamics simulation of 200ns for each protein-ligand complex was performed in GROMACS. The identified ligands are found to impart higher rigidity and stability to the protein-ligand complexes. Furthermore, the results were validated by performing the principal component analysis (PCA), and calculation of binding free energy using the Molecular Mechanics Poisson-Boltzmann Surface Area (MMPBSA) approach. The identified ligands occupy smaller phase space, form stable clusters and exhibit stronger non-bonded interactions. Thus, our findings can be useful for the development of promising anti-fibrinolytic agents.

**Graphical Abstract:** 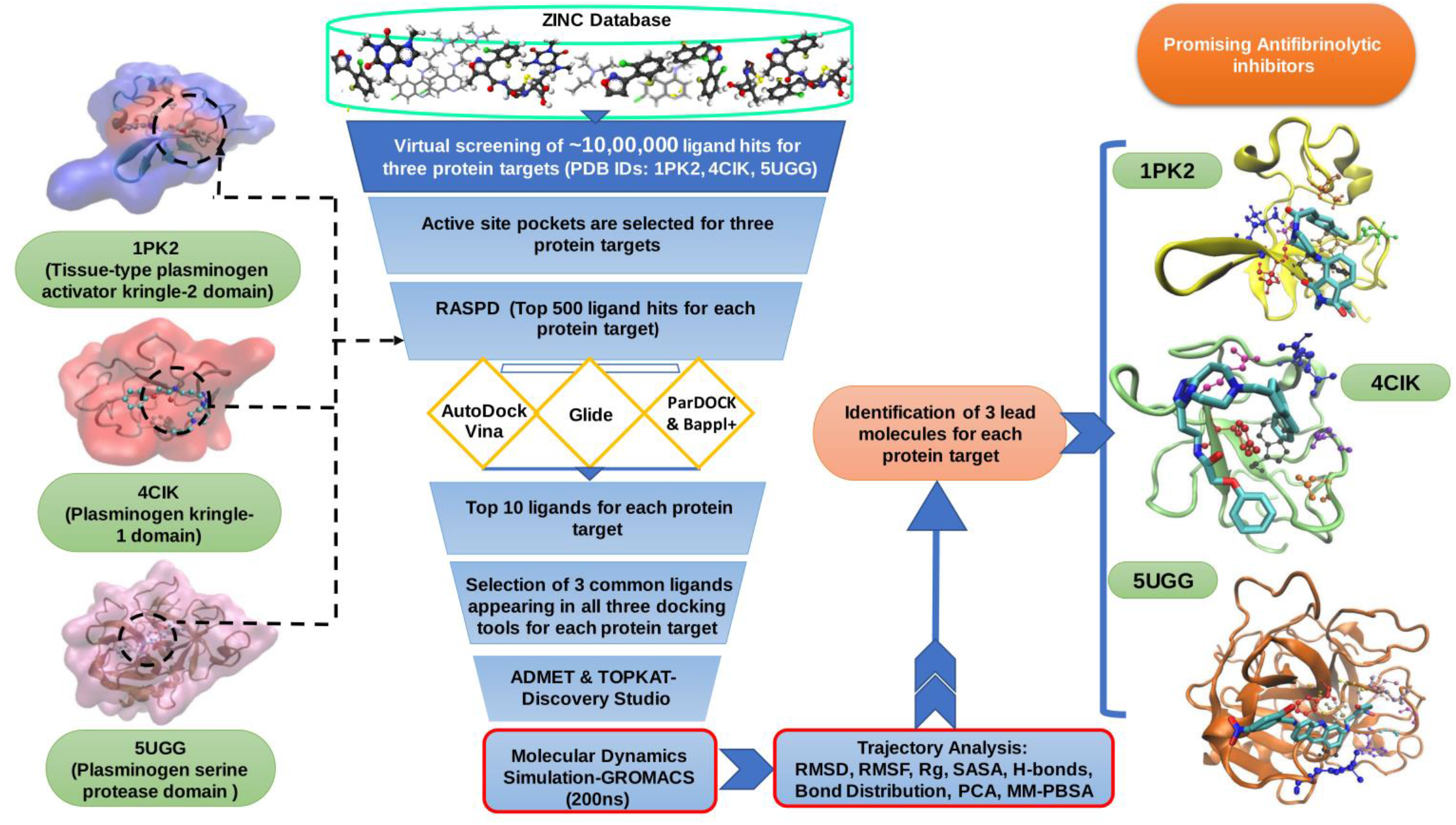

## 1. Introduction

Fibrinolysis is a process of removal (lysing) of clots formed due to the triggering of the hemostatic pathway, in conditions such as pathological thrombosis and vascular occlusion (Chapin et al., 2015). The principal mediator of fibrinolysis is plasminogen, which is transformed into plasmin by the plasminogen activators (Tissue-type plasminogen activator and Urokinase plasminogen activator) (Harpel et al., 1985). Tissue-type plasminogen activator (tPA or PLAT) is a serine protease found on the surface of endothelial cells that plays an important role in degrading polymerized fibrin by hydrolyzing the Arg560-Va1561 peptide bond in plasminogen (Plg) (Fears R., 1989). t-PA consists of six well-defined structural modules: a finger domain; an epidermal growth factor domain; two kringle domains and a serine protease two-domain unit with trypsin-like specificity (Novokhatny et al., 1991). Because of the presence of a lysine binding site in the above-mentioned two kringle domains and its stimulatory effect on Plg activation by fibrin, the scientific community has shown significant interest in the kringle domain (Verheijen et al., 1986). The zymogen protease Plg and its active form plasmin (Plm), carry out important functions in blood clot disintegration apart from other cellular processes such as bacterial pathogenesis, tissue remodeling, and cell migration (Korhonen et al., 2013). Plg comprises a serine protease (SP) domain, a Pan-apple (PAp) domain, and 5 kringle (KR 1-5) domains. These domains are ordered in a closed configuration that is sustained through interdomain interactions facilitated by the canonical lysine-binding sites (LBSs) of the KR domains. The binding of Plg to lysine-containing substrates or receptors is also facilitated by the LBSs of the KRs. Post binding, Plg takes an open configuration that is triggered by tissue plasminogen activators (tPAs) or urokinase plasminogen activators (uPAs) (Lijnen et al., 1980). To regulate this binding process, α_2_-antiplasmin and α_2_-macroglobulin block the release of activated Plm from the clot or the cell surface (Ayón-Núñez et al., 2018). Irregular balance of Plg/Plm system and immoderate fibrinolytic activity leads to fatal hemorrhagic disorders, thrombotic vascular injury, and severe complications during general surgery or major trauma (Colomina et al., 2021). Presently, the Plm inhibitors aprotinin and tranexamic acid (TXA) are clinically used to treat severe hemorrhages by blocking new entry to target substrates and are the most extensively used inhibitor for the Plg/Plm system (McCormack et al., 2012). TXA is a reasonably well-tolerated lysine analog that binds to the LBSs, culminating in the development of open Plg (Nishida et al., 2017). Post binding, TXA obstructs Plm activation by stopping it from binding to substrates and/or receptors. However, one recent study has shown that TXA may not block the activity of Plm which has already been formed leading to dangerous complications in patients suffering from extreme blood loss (Roberts et al., 2013). Hence, there is a requirement to synthesize specific inhibitors which can attach to the protease active site of Plg. Cross-reaction of Plm inhibitors with other plasma serine proteases also challenges their development because the reactions generate multiple side effects like headaches, nasal symptoms, or back, abdominal, and muscle pain. In some cases of treatment with TXA, seizures were reported as a side effect which in turn were traced to gamma-aminobutyric acid antagonistic activity (GABAa) (Hunter & Young, 2011). Also, the gastrointestinal side effects associated with a high dose of TXA can be attributed to GABAa (Ockerman et al., 2021). Based on the findings of these studies, the ideal novel oral fibrinolysis inhibitor is expected to work on a mechanism of action similar to that of TXA, having suitable dosing and higher selectivity over GABAa. Of late, some new encouraging drugs have been discovered like 4-PIOL and PS-112 (TXA-derived active site inhibitors) but, to date, their detailed profiling and testing have not been performed (Mortensen et al., 2019; Law et al., 2017). Even though considerable research has been undertaken, clinically safe fibrinolysis inhibitors have not reached the market. Fibrinolysis can be suppressed by targeting each of the three important protein domains: the kringle-2 domain of tissue plasminogen activator, the kringle-1 domain of plasminogen, and the serine protease domain of plasminogen. Previous studies reported that the catalytically active domain is constituted by the serine protease domain while the kringle domains are crucial for protein-protein interactions such as binding to fibrin (Wiman & Collen, 1978). Therefore, we have selected the above-mentioned three individual protein domains as our respective targets (Van Zonneveld et al., 1986; Castellino FJ et al., 2005; Keragala &Medcalf, 2021). Herein, we attempt to find potential inhibitors having minimum side effects with maximum specificity and efficacy using *in silico* approaches. In this present study, combined approaches of structure-based virtual screening and molecular docking were employed to discern prospective hits from the ligand database (Lionta et al., 2014). Thereafter, ADMET (Absorption, Distribution, Metabolism, Excretion, and Toxicity) properties of these identified ligands were evaluated and a molecular dynamics simulation of 200ns was carried out for each protein-ligand complex. Furthermore, principal component analysis (PCA) was performed and binding free energy was calculated using the Molecular Mechanics Poisson-Boltzmann Surface Area (MMPBSA) approach for all the protein-ligand complexes. The identified lead compounds are expected to contribute to the development of potent anti-fibrinolytic agents.

## 2. Methods

### 2.1. Protein Preparation

The crystal structures of the kringle-2 domain of tissue plasminogen activator, the kringle-1 domain of plasminogen, and the serine protease domain of plasminogen with PDB IDs: 1PK2, 4CIK, 5UGG were retrieved from the protein data bank (PDB) respectively (Byeon IJ & Llinás M, 1991; Cheng L et al., 2014; Berman et al, 2000). All three protein structures were analyzed in the pymol and discovery Studio 3.5 package (Accelrys, San Diego, CA, USA), and the missing residues in the structures were added using Modeller (The PyMOL Molecular Graphics System, Version 2.0 Schrödinger, LLC; Webb & Sali, 2016; Systemes, D. BIOVIA. “Discovery Studio Modeling Environment.” Dassault Systèmes, San Diego, 2016). The co-crystallized ligands with IDs: ACA, XO3, 89M were extracted from protein crystal structures 1PK2, 4CIK, and 5UGG respectively using discovery Studio 3.5. The water molecules were eliminated, and protein preparation was done using the default settings of discovery Studio 3.5. Further, the protein structures were optimized using CHARMM (Chemistry at Harvard Macromolecular Mechanics) force field, and energy minimization was done by a smart minimizer algorithm incorporating conjugate gradient energy protocol (Brooks et al., 2009). Subsequently, these protein structures were selected as targets for virtual screening.

### 2.2. Structure-based virtual screening

The library of one million ligands was retrieved from the ZINC 15 database (Sterling & Irwin, 2005). Initially, these one million ligands were screened individually against 1PK2, 4CIK, and 5UGG protein targets respectively using RASPD (Mukherjee & Jayaram, 2013). This tool rapidly identifies hit molecules against the protein targets based on their physicochemical properties and calculates the binding energy. Subsequently, the hits were further evaluated based on Lipinski parameters such as hydrogen bond acceptors & donors, molar refractivity, ADMET, Wiener index, and volume of functional groups. The residues which were bound to the co-crystallized ligands were selected as active site pockets of the target proteins for virtual screening. From RASPD, the top 1500 hits (500 for each protein target) were retrieved for further analysis.

### 2.3 Molecular Docking

The top 500 hits filtered out from virtual screening (1500 total) were subjected to molecular docking against the three protein targets 1PK2, 4CIK, and 5UGG using three docking tools such as Schrödinger-Glide, Autodock Vina and ParDOCK/BAPPL+ respectively (Trott & Olson, 2010; Glide version 6.2. (2019). New York, NY: Schrodinger, LLC; Gupta et al., 2007; Soni et al., 2020)

#### 2.3.1. Glide (Grid-based ligand docking with energetic)

Firstly, the ligands were prepared using the LigPrep module of Schrödinger Maestro version 2.9, 2019 so that the ligands could be supplied to glide for docking in a state as they would occur in a protein-ligand complex (LigPrep, version 2.9. (2013). New York, NY: Schrodinger, LLC). LigPrep uses the Optimized Potential Liquid Simulation (OPLS) 2005 force field and consists of a series of steps to optimize the structures. By default, LigPrep adds hydrogens, removes unwanted molecules, and minimizes the ligand structure. Thereafter, docking was performed using the software package Glide version 2.9, 2019. Protein structures were prepared in the Protein Preparation Wizard of Schrödinger in which proteins were refined by adding missing hydrogens and assigning proper bond orders. A grid file was generated using the Receptor Grid Generation protocol on these minimized structures, such that the grid points were adjusted to the active site pocket in X, Y, and Z dimensions with 25 × 25 × 25 points, and the ligands were allowed to move freely. We have used Glide docking modules namely simple precision (SP) and extra precision (XP) for docking calculations (Friesner et al., 2006). The conformations obtained from SP were used as input for XP to generate multiple binding poses. The binding affinity was computed based on the glide scoring function for all three target proteins i.e., 1PK2, 4CIK, and 5UGG respectively. Thereafter, the top ten ligands were ranked according to their binding affinity using GScore (GlideScore) scoring function which is a combination of Coulombic energies, Van der Waals energies, and the internal strain of the ligand (Halgren et al., 2004).

#### 2.3.2. AutoDock Vina

As mentioned earlier, 1500 hits (500 hits per protein) were processed and subjected to docking analysis in AutoDock Vina. ADT tool was used to load the target protein structure and convert proteins to PDBQT format. Subsequently, the ligand hits were also converted into PDBQT format in Open Babel (O’Boyle et al., 2011). The hydrogen atoms and water molecules from the protein structure were removed and polar hydrogen atoms were added. Thereafter, Kollman charges were incorporated into the target protein structures (Kollman et al., 2000). The auto-grid box was set across X-Y-Z directions, i.e., 23×20×18 points for tissue-type plasminogen activator kringle-2 domain (PDB ID: 1PK2), 18×20×18 points for plasminogen kringle-1 domain (PDB ID: 4CIK), and 29×28×28 points for serine protease domain of plasminogen (PDB ID: 5UGG) respectively. The grid was set up to encompass the active site pocket, with a spacing of 1 Å. The ligand hits were ranked according to their binding energies and finally, the best-performing top ten ligands for each protein target were selected.

#### 2.3.3. ParDOCK & Bappl+

The same set of 1500 hits (500 hits per protein) was screened by the Monte Carlo method involving an all-atom energy-based system using default parameters of ParDOCK software, hosted in the Sanjeevani server (Jayaram et al., 2012). The ligands were positioned optimally around the active site of the protein target and ranked based on their interacting energies. Subsequently, the scoring function of all the aforementioned ligands was derived using the Bappl+ tool which utilizes the Random Forest algorithm scoring function (Li et al., 2015). In Bappl+ the default parameters were set and formal charges were assigned for each ligand. Based on these settings, the binding affinities of ligands were computed and the top ten compounds were ranked accordingly.

### 2.4. ADMET and TOPKAT

All the top ten identified ligands obtained from Glide, Autodock Vina, ParDOCK, and Bappl+ tools were visually inspected in Pymol and Schrödinger maestro. Based on the docking scores, out of these ten identified ligands, the top three were selected for ADMET and TOPKAT (Toxicity Prediction by Computer Assisted Technology) analysis for each protein target. ADMET properties such as CYP2D6 & plasma protein inhibition, intestinal adsorption, aqueous solubility, hepatotoxicity, blood-brain barrier level were evaluated using the Discovery studio 3.5 ADMET tool kit. Furthermore, Lipinski’s rule of five was also checked for these ligands (Lipinski, 2004). Thereafter, the toxicity of each ligand was estimated using the TOPKAT tool, which contains a robust Quantitative Structure-Toxicity Relationship (QSTR) modeling system to predict the accurate toxicity endpoints (Prival, 2004). The toxicity properties such as rodent carcinogenicity, Ames mutagenicity, skin irritation, and developmental toxicity potential were also investigated.

### 2.5. Molecular Dynamics Simulation (MD)

The top three ranked ligands for each protein target were subjected to MD simulation using GROMACS (Groingen Machine for Chemical Simulations) 2019 suite implementing GROMOS (54a7) force field (Prival et al., 2001; Pronk et al.,2013; Huang et al., 2011). The protein-ligand complex was solvated in a cubic box of 1 nm using periodic boundary conditions and a Simple point-charge water model (SPCE) (Lombardero et al., 1999). The Na+ and Cl- ions were added to neutralize the system and maintain the concentration of 0.15 mol/L. The PRODRG server was used to generate the ligand parameters and topology (Schüttelkopf & Van Aalten, 2004). After that, internal constraints of the protein-ligand complex were relaxed by 2000 steps of steepest descent energy minimization with a max force constraint of 1000 KJ/mol, leading to restraining positions of all heavy atoms. Before MD simulations, the systems were heated using a V-rescale thermostat to attain the temperature of 310 K with 0.1ps as the constant of coupling and achieved equilibration in NVT (Number of atoms, Volume of the system, and Temperature of the system). Then solvent density was sustained using a Parrinello-Rahman barostat with the pressure of 1 bar, coupling constant of 0.1 ps, and temperature of 310 K to obtain equilibration in NPT (Number of atoms in the system, the pressure of the system, and temperature of the system) by gradually discharging the restraint on heavy atoms step by step (Parrinello & Rahman, 1981). Finally, an MD simulation was performed for the equilibrated structures for 5 ns with an integration time step of 2fs. The electrostatic interactions of long-range were implemented using particle-mesh Ewald sum with a cutoff of 1.0 nm (Essmann et al., 1995). During simulation, the LINCS algorithm was used to constrain all bond lengths, and the SETTLE algorithm was used for restraining water molecules (Hess et al., 1997; Miyamoto & Kollman, 1992). The resultant structure from the NPT equilibration phase was employed for the final production run in the NPT ensemble for 200 ns simulation time. Finally, the trajectory analysis such as RMSD, RMSF, Rg, SASA, H-bonds, PCA of protein-ligand complexes was performed using gromacs utilities and plotted in xmgrace (Turner, 2005).

### 2.6. Principal Component Analysis

PCA is a statistical method that is applied to reduce data complexity and is useful for analyzing the large-scale motion of proteins (David & Jacobs, 2014). Generally, internal motions of every protein molecule are ingrained and are required for their proper biological functioning like conformational changes to various biological environments, binding to substrate, etc. Due to problems in interpreting these internal motions of the protein, PCA is utilized to minimize the large dimensions of the data set to identify the prominent principal components (PCs). These PCs represent the major contributors responsible for elucidating crucial information about the dynamic changes of the protein. In PCA, a covariance matrix was constructed from trajectory data after eliminating unwanted motions (translational and rotational). This was followed by diagonalization of the covariance matrix using the gmx_covar tool which is a part of the Gromacs module. By diagonalization of covariance matrices containing backbone C alpha atoms data, the eigenvector and eigenvalues were obtained which corresponds to the change in protein trajectory throughout the simulation time. The *gmx anaeig* tool of Gromacs was used to analyze and plot the trajectories of the backbone C alpha atoms of all the systems. In our study, the first two projections (PC 1 and PC 2) with the highest eigenvalues were taken into consideration as they are responsible for 80–90% of the collective motions of the C alpha backbone atoms.

### 2.7. Free energy calculation

To evaluate the binding free energy (ΔG) of protein-ligand complexes, the Molecular Mechanics Poisson-Boltzmann Surface Area (MM-PBSA) method was employed (Genheden & Ryde,2015). We computed ΔG for the last 50 ns of the production run using the *g_mmpbsa* tool of the Gromacs module (Kumari & Kumar, 2014). Binding free energy was calculated for the identified ligands by estimating the bound and unbound state differences with the protein targets.

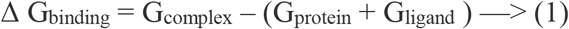

In equation (1), the free energies of protein and ligands are indicated by G _protein_ and G _ligand_. The free energy of the protein-ligand complex is represented by the G _complex_

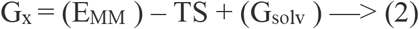

Similarly, equation (2) is used to calculate the bound or unbound state’s free energy. The x indicates the unbound states or protein-ligand complex. E _MM_ is used to calculate the average molecular mechanics energy and TS indicates the entropic contribution. The solvation free energy is indicated by G _solv_

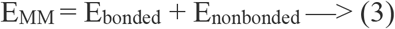

Equation (3) shows that bonded and non-bonded (i.e. vander Waal’s & electrostatic) interactions were considered for calculating the molecular mechanics energy (E_MM_) for protein-ligand interaction.

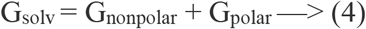

The linearized Poisson Boltzmann equation is represented by G_solv,_ where The polar component G_polar_ refers to the amount of energy needed to move a solution from a continuous media with a low dielectric constant (ε= 1) to one with water’s dielectric constant (ε= 80).

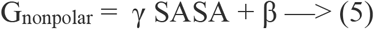

Where, γ = 0. 0227 kJ mol^-1^ Å^-2^ and β = 0 kJ mol^-1^ (Brown & Muchmore, 2009). A probe with a radius of 1.4 is used to define the dielectric boundary. The nonpolar contribution G_nonpolar_ is taken into consideration for estimating the solvent accessible surface area (SASA). It was assumed that as all ligands are binding to one distinct protein target, therefore these ligands will contribute similar entropic energy. Hence, in our analysis complicated entropic contribution was not taken into account and was removed at the time of calculation.

## 3. Results and discussion

### 3.1 Virtual screening (VS)

VS is the most widely used technique in the drug discovery process. The complete flow chart of our current study is represented in Figure 1. We have selected three protein targets which known to play a crucial role in the fibrinolysis process i.e., tissue-type plasminogen activator kringle-2 domain (PDB ID: 1PK2), plasminogen kringle-1 domain (PDB ID: 4CIK), and serine protease domain of plasminogen (PDB ID: 5UGG). From the crystallographic structures of protein i.e., 1PK2, 4CIK, and 5UGG, the co-crystallized ligands (ACA, XO3, and 89M) were removed from the protein complex, and the residues found interacting with these ligands were selected as active site pockets. We have used the structure-based drug design approach to identify the lead compounds which could bind to the specific pocket region accordingly (Anderson AC, 2003). One million molecules were retrieved from the ZINC database and post-screening of these compounds in the RASPD tool we obtained 500 hits for each protein target (1500 hits total). These 1500 compounds (500 per protein) were optimized in the Discovery studio and molecular docking was performed by using three different tools accordingly.

**Figure 1:**
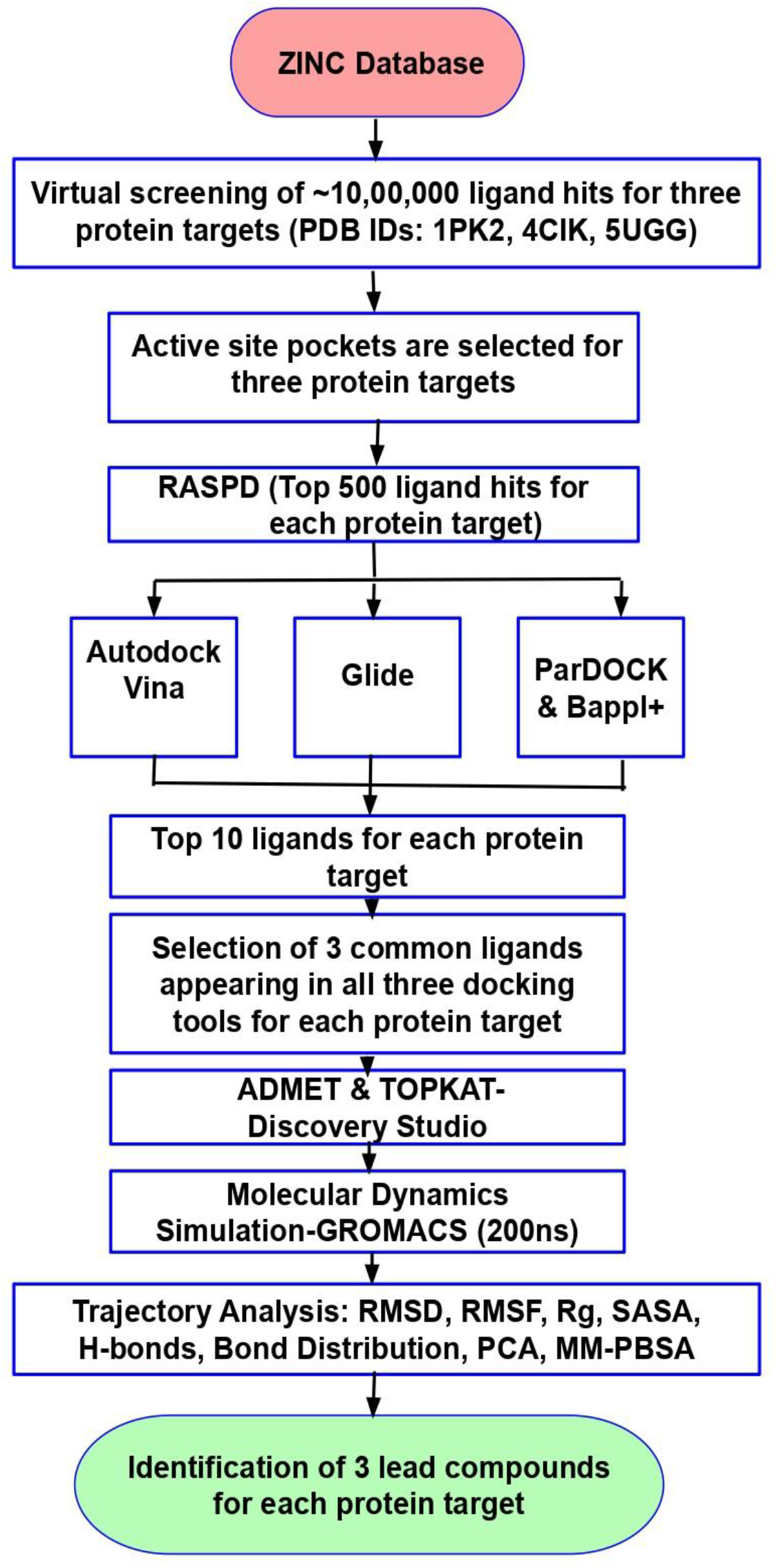
Flow chart of the virtual screening process using combined molecular docking and molecular dynamics simulation approaches.

### 3.2 Comparative Molecular Docking

To understand the interaction profile of 1500 ligands (500 per protein) with the protein targets, we performed a comparative molecular docking analysis using three different tools, i.e., Auto dock Vina, Glide, and ParDOCK/BAPPL+. The 500 hits were individually docked to the active site pocket of 1PK2, 4CIK, and 5UGG respectively. The key active site residues of the 1PK2 protein target include VAL40, ASP63, TRP69, ARG71, LEU78, THR79, TRP80. Similarly, the binding site of the 4CIK protein target includes ARG35, ASP55, ASP57, TRP62, TYR64, ARG71, TYR72 active site residues. Likewise, for the 5UGG protein target, the residues comprising the docking site include PHE587, CYS588, HIS603, LYS607, SER736, CYS737, GLN738, GLY739, SER741, THR759, TRP761, and GLY762. Based on the binding energies, the top 10 hits for each target protein were ranked based on the scores obtained from the three docking tools: Autodock vina, ParDOCK/BAPPL+, and Glide (Supplementary Table ST1, ST2, and ST3). Among these 10 hits, we narrowed it down to the top three ligands by selecting the common hits found in all three docking tools. The docking results obtained from docking all the nine identified ligands (P27, P67, P76, C19, C90, C97 U19, U94, U97) and the reference ligands (ACA, XO3, and 89M) with their respective protein targets (1PK2, 4CIK, and 5UGG) are provided in Table 1. Detailed Glide SP and XP scores of the top three ligands hit against each protein target are also provided in (Supplementary Table ST4). The interacting residues which include both the active site and novel (other than the active site) residues of the proteins (1PK2, 4CIK, and 5UGG) with their respective ligands (P27, P67, P76, C19, C90, C97 U19, U94, U97) 1PK2 are provided in Table 1. The 2D structures of the nine identified compounds (three for each protein target) are provided in Figure 2 [(A)-(I)]. For comparative docking studies of the 1PK2 protein target, the ligand ACA was selected as the reference ligand. Its binding energy was found to be -3.9 kcal/mol and -4.9 kcal/mol in Autodock Vina and ParDOCK/BAPPL+ respectively. Similarly, the Glide score obtained for the reference ligand was -7.25 kcal/mol. Simultaneously, we conducted docking studies for each of the top three ligands i.e., ZINC12211330 (P27), ZINC00718625 (P67), ZINC09970930 (P76) against the 1PK2 protein target. The identified ligands P67, P27, P76 exhibited binding energy of -7.6, -8.3, -8.9 kcal/mol in Autodock Vina and -8.84, -9.60, -9.63 kcal/mol. in ParDOCK/ BAPPL+ respectively. Similarly, the Glide score obtained for P67, P27 & P76 was -8.6, -9.31, -9.42 kcal/mol respectively. As shown, the interacting residues of 1PK2 with P27 are LYS39, VAL40, TYR41, ASP63, TRP69, HIS71, ARG77, LEU78, THR79, TRP80, GLU81, TYR82. Likewise, the interacting residues of 1PK2 with P67 are GLU23, LYS39, VAL40, TYR41, HIS71, ASP63, TRP69, ARG77, LEU78, TRP80, GLU81, TYR82. Similarly, the interacting residues of 1PK2 with P76 are LYS39, VAL40, TYR41, TRP69, HIS71, ARG77, LEU78, THR79, TRP80, GLU81, TYR82. Docking studies for each of the top three ligands i.e., ZINC02060288 (C19), ZINC12455413 (C90), ZINC14888376 (C97) were performed against the 4CIK protein target. The identified ligands C90, C19, C97 exhibited binding energy of -8.2, -8.5, -8.8 kcal/mol in Autodock Vina and -8.90, -9.0, -9.47 kcal/mol in ParDOCK/ BAPPL+ respectively. Similarly, the Glide score obtained for C90, C19, C97 was -8.75, -8.92, -9.3 kcal/mol respectively. The reference ligand XO3 exhibited binding energy of -7.4 kcal/mol in Autodock Vina and - 6.66 kcal/mol in ParDOCK/BAPPL+ and -7.25 in Glide respectively. As shown, the interacting residues of 4CIK with C19 are ARG35, ASP55, ASP57, PRO58, GLN59, TRP62, TYR64, ARG71, TYR72, TYR74. Likewise, the interacting residues of 4CIK with C90 are ARG35, ASP55, ASP57, GLN59, TRP62, GLU69, LYS70, ARG71, TYR72, ASP73, TYR74. Similarly, the interacting residues of 4CIK with C97 are ARG35, PHE36, ASP55, ASP57, PRO58, GLN59, TRP62, TYR64, ARG71, TYR72, TYR74. In the same way, docking studies for each of the top three ligands i.e., ZINC12576410 (U19), ZINC04557820 (U94), ZINC11839443 (U97) were conducted against the 5UGG protein target. The identified ligands U19, U94 and U97 exhibited binding energy of -10.9, -11.0, -11.4 kcal/mol in Autodock Vina and -7.9, -8.02, -8.33 kcal/mol in ParDOCK/ BAPPL+ respectively. Similarly, the Glide score obtained for U19, U94, and U97 was -8.87, -9.41, -9.6 kcal/mol respectively. The reference ligand 89M exhibited binding energy of -9 kcal/mol in Autodock Vina and -5.9 kcal/mol in ParDOCK/BAPPL+ and -7.5 in Glide respectively. As shown, the interacting residues of 5UGG with U19 are HID63, CYS64, LEU65, GLU66, LYS67, SER68, ARG70, SER72, SER73, TYR74, GLN198, SER201. Likewise, the interacting residues of 5UGG with U94 are SER68, LYS67, TYR74, GLU66, LEU65, CYS64, HID63, CYS197, SER196, ASP195, VAL233, GLY232, CYS225, GLY224, LEU223, GLY222, TRP221, SER220, SER201. Similarly, the interacting residues of 5UGG with U97 are ALA62, HID63, CYS64, LEU65, GLU66, LYS67, SER68, TYR74, SER73, CYS197, SER196, ASP195, GLY232, VAL233, SER220, TRP221, GLY222, LEU223, GLY224, SER201. Overall, from the docking results, it can be construed that in comparison to reference ligands ACA, XO3, and 89M, the identified ligands P27, P67, P76, C19, C90, C97 U19, U94, U97 exhibited higher binding energy and stronger affinity towards their respective protein targets 1PK2, 4CIK, and 5UGG. Apart from the active site residues of the target proteins, no other residues are found to interact with the reference ligand ACA, XO3, and 89M. Therefore, it can be ascertained that the identified ligands form more favorable interactions with their respective protein targets, thus enhancing stronger and stable binding. The 2D interaction images of all the top nine ligands P27, P67, P76, C19, C90, C97 U19, U94, U97 complexed with their respective protein targets 1PK2, 4CIK, and 5UGG are also provided in Figure 3 [(A)-(I)]. It is noteworthy to mention that out of the nine identified ligands, P76, C97, and U97 are the top three ligands that bind with greater affinity to their respective targets 1PK2, 4CIK, and 5UGG as implied from the docking scores obtained from Autodock vina, ParDOCK/BAPPL+ and Glide (Table1). Post docking, ADMET & TOPKAT properties such as solubility, intestinal adsorption, etc. of the identified ligands and the reference ligands were investigated. As provided in the supplementary table ST5 and ST6, these ligands were found non-hepatotoxic, have the least development toxicity and don’t inhibit the cytochrome 4502D6 (CYP2D6) enzyme. The discovered ligands were also found not to bind to Plasma Protein (PP) and to be free of carcinogenicity, mutagenicity, and skin irritation in rodents.

**Table 1:**
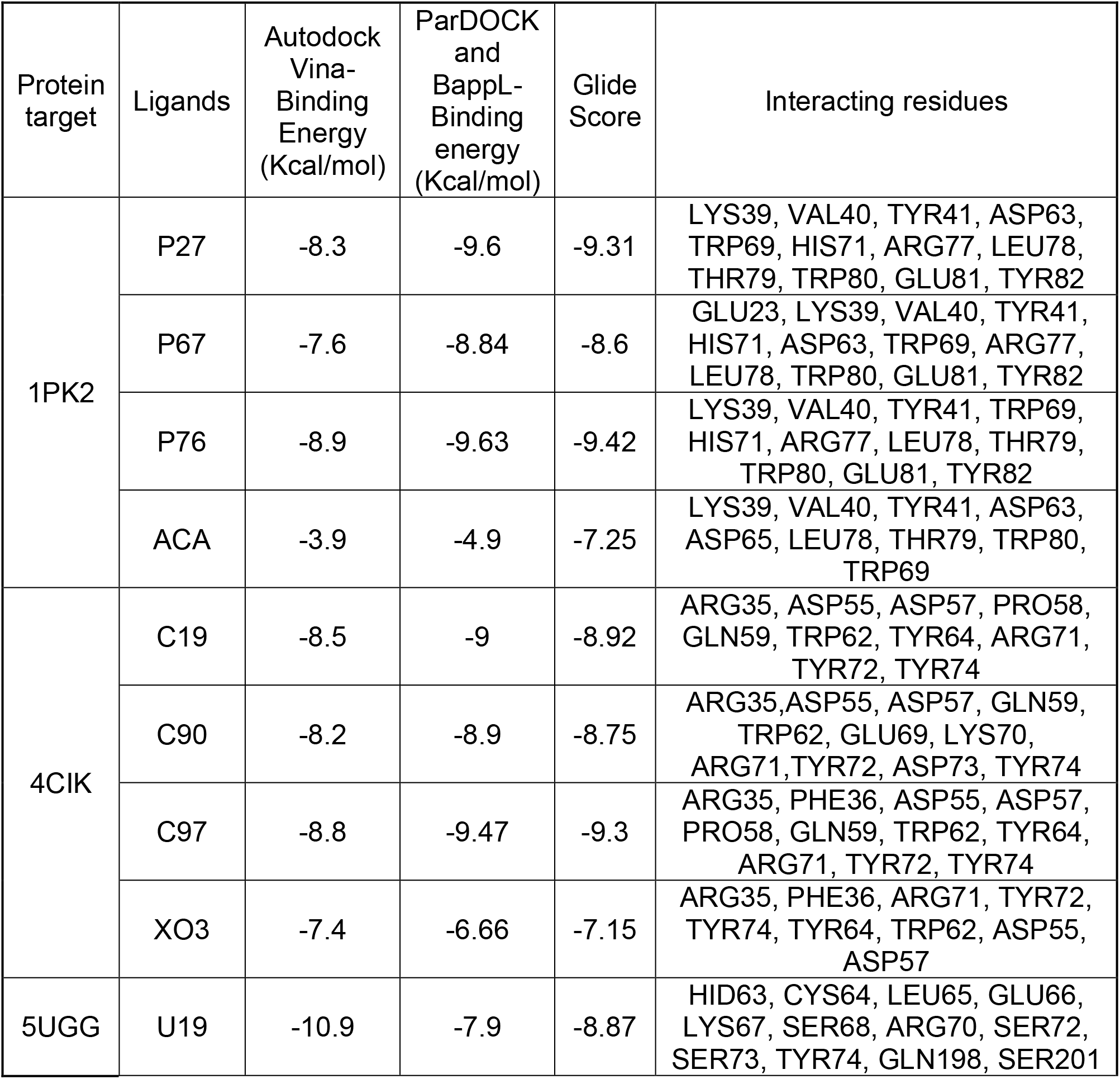
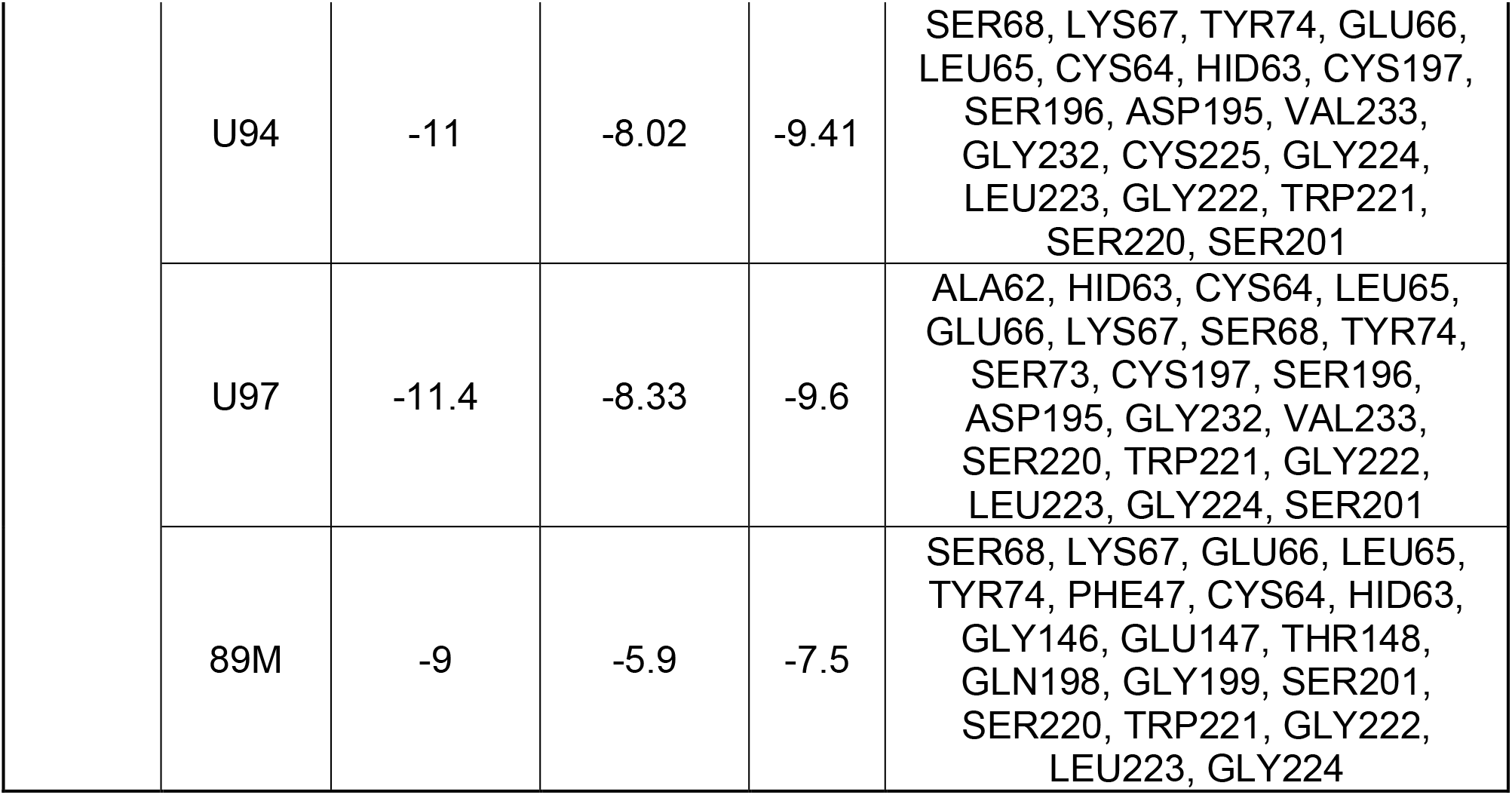
Molecular docking results of top nine identified ligands (P27, P67, P76, C19, C90, C97 U19, U94, U97) and the reference ligands (ACA, XO3, and 89M) bound with their respective protein targets (1PK2, 4CIKand 5UGG) using three docking tools: Autodock vina, ParDOCK/BAPPL+, and Glide respectively.

**Figure 2:**
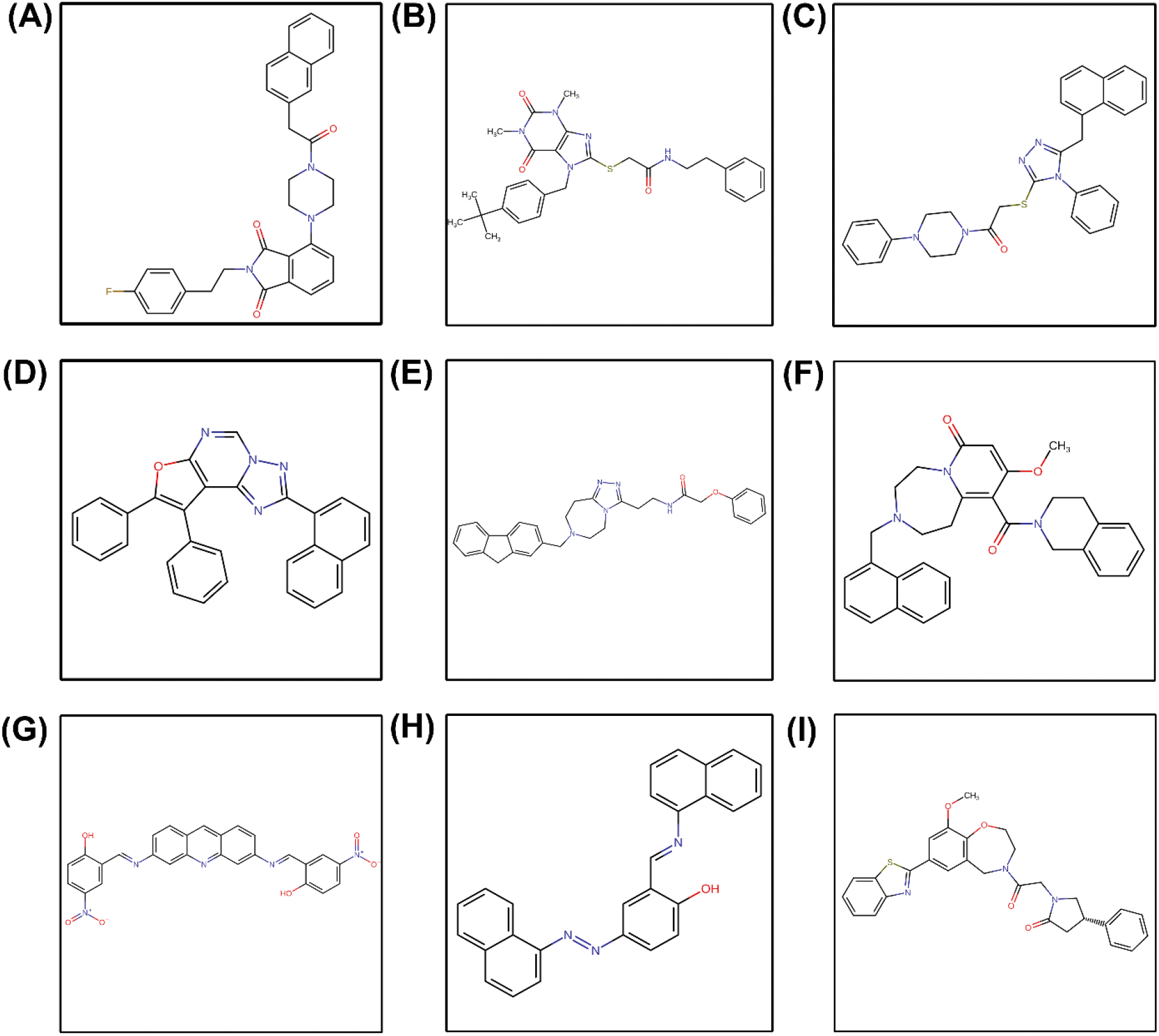
The 2D structures of the lead identified compounds (A) ZINC12211330 (P27), (B) ZINC00718625 (P67), (C) ZINC09970930 (P76), (D) ZINC02060288 (C19), (E) ZINC12455413 (C90), (F) ZINC14888376 (C97), (G) ZINC12576410 (U19), (H) ZINC04557820 (U94), (I) ZINC11839443 (U97)

**Figure 3:**
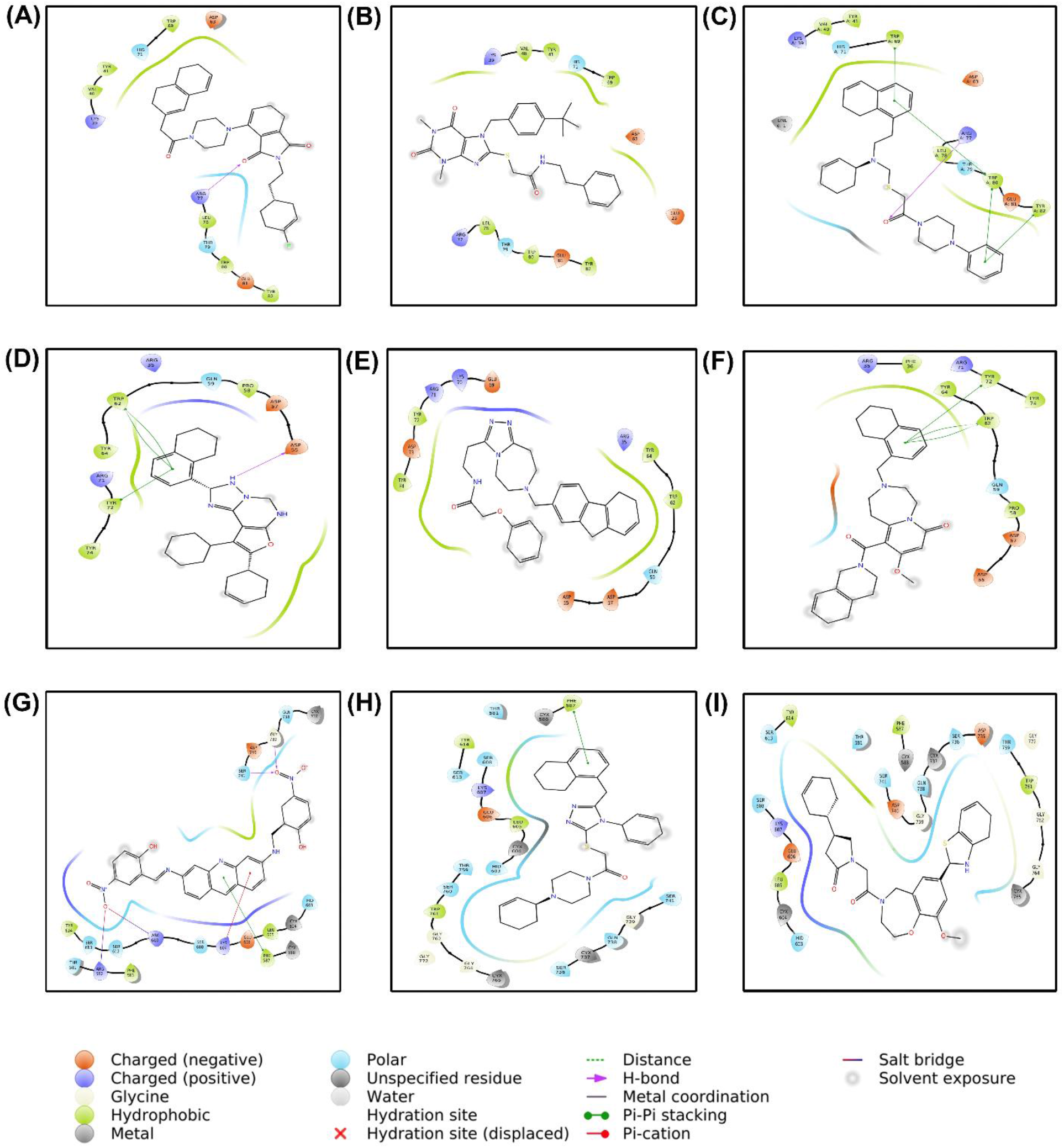
The 2D interaction profile of lead molecules into the binding site of their respective protein targets 1PK2, 4CIK and 5UGG (A) ZINC12211330 (P27), (B) ZINC00718625 (P67), (C) ZINC09970930 (P76), (D) ZINC02060288 (C19), (E) ZINC12455413 (C90), (F) ZINC14888376 (C97), (G) ZINC12576410 (U19), (H) ZINC04557820 (U94), (I) ZINC11839443 (U97)

### 3.3 Molecular Dynamics Simulations

Molecular Dynamics Simulation (MDS) plays a significant role in studying conformational changes of protein-ligand complexes and provides valuable insights into the prediction and identification of protein-ligand interactions (Durrant & McCammon, 2011). We have performed MD simulations of each system (apo protein, protein-bound with reference ligands, and protein-bound with identified ligands) for 200ns. Thereafter, we performed trajectory analysis such as Root-mean-square-deviation (RMSD), Root-mean-square fluctuation (RMSF), Radius of gyration (Rg), number of hydrogen bonds, bond distribution, Principal component analysis (PCA), and binding free-energy calculation (MMPBSA) for all the systems (apo proteins, protein-bound with reference ligand complexes, and protein-bound with identified ligand complexes).

#### 3.3.1 Root-mean-square deviation analysis

To elucidate the influence of ligands on the conformational stability of the protein, we have analyzed the root-mean-square deviation (RMSD) of backbone atoms of all the systems using the standard *g_rms* function of GROMACS for an overall time of 200 ns simulation run (Figure 4A-4C). As provided in supplementary table ST7, 1PK2 protein docked with the reference ligand ACA and the identified ligands P27, P67, P76 shows an average deviation of 0.32± 0.04 nm, 0.45± 0.05 nm, 0.36± 0.04 nm, 0.4± 0.03 nm respectively. The 1PK2 APO protein exhibits an average deviation of 0.44± 0.05 nm. From the RMSD graphs, it can be inferred that all the systems are equilibrated and have converged within a range of 0.2-0.5nm (Figure 4A). Similarly, 4CIK protein docked with ligands X03 (reference ligand), C19, C90, C97 shows an average deviation of 0.29± 0.03 nm, 0.29± 0.03 nm, 0.3± 0.03 nm, 0.34± 0.04 nm respectively. The 4CIK APO protein exhibits an average deviation of 0.34± 0.06 nm. This suggests that all the systems are equilibrated and have converged within a range of 0.2 - 0.4 nm (Figure 4B, supplementary table ST7). Likewise, 5UGG protein docked with ligands 89M (reference ligand), U19, U94, U97 exhibited an average deviation of 0.24± 0.02 nm, 0.29± 0.02 nm, 0.3± 0.02 nm, 0.31± 0.02 nm respectively. The 5UGG APO protein shows deviations of 0.24± 0.02 nm. This implies that all the systems that have converged within a range of 0.2-0.35 nm have attained equilibration (Figure 4C, supplementary table ST7). Although the RMSD profile shows an insignificant difference in the average value and minor fluctuations throughout, convergence is observed which indicates that our simulations have achieved stable trajectories. Hence, other conformational dynamics analysis is required to deduce further conclusions.

**Figure 4:**
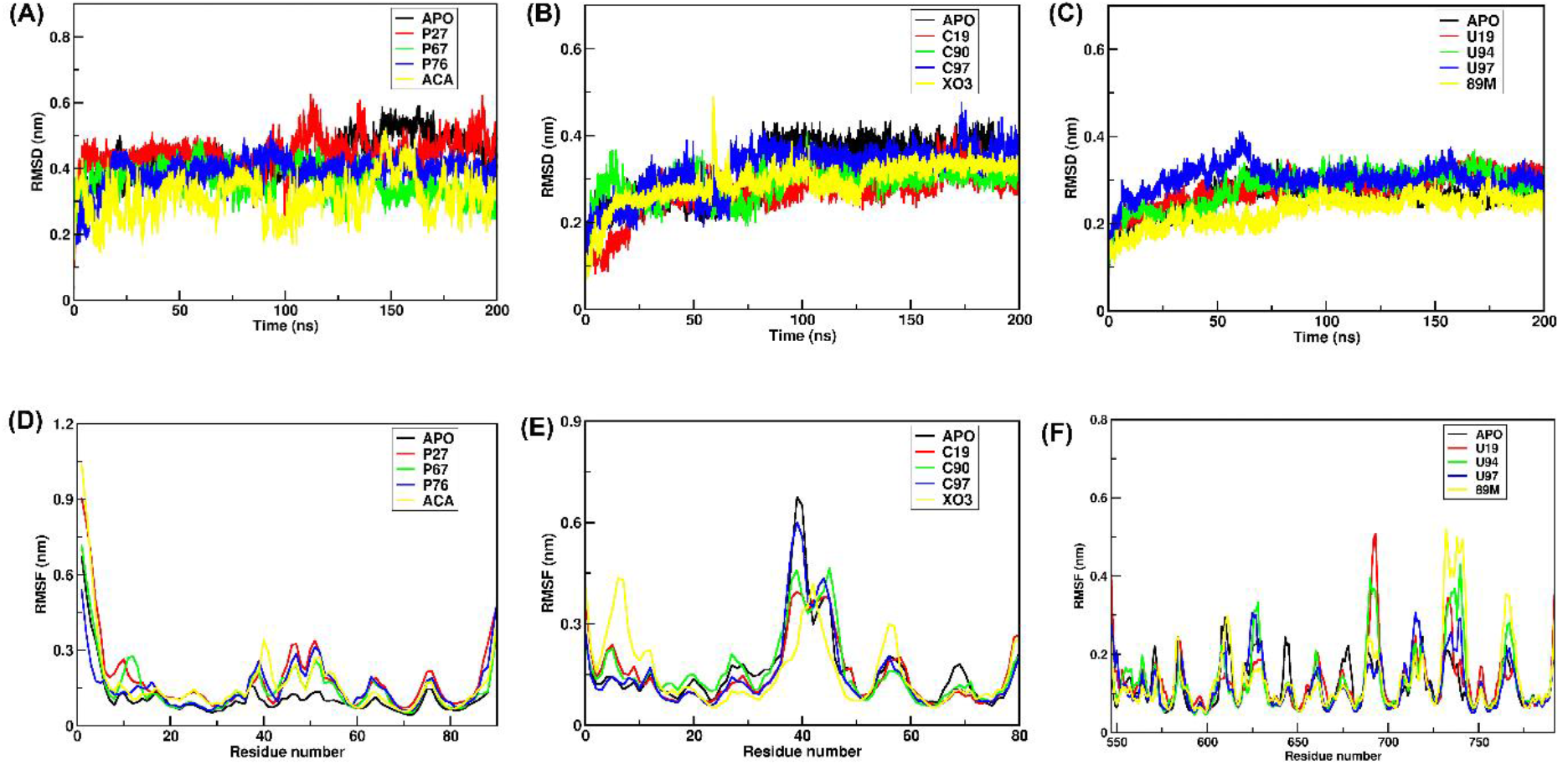
(A) RMSD profile of 1PK2 protein-ligand complexes (top three identified ligands P27, P67, P76, and reference ligand ACA) along with APO. (B) RMSD profile of 4CIK protein-ligand complexes (top three identified ligands C19, C90, C97and reference ligand XO3) along with APO. (C) RMSD profile of 5UGG protein-ligand complexes (top three identified ligands U19, U94, U97, and reference ligand 89m) along with APO. (D) RMSF profile of 1PK2 protein-ligand complexes (top three identified ligands P27, P67, P76, and reference ligand ACA) along with APO. (E) RMSF profile of 4CIK protein-ligand complexes (top three identified ligands C19, C90, C97and reference ligand XO3) along with APO. (F) RMSF profile of 5UGG protein-ligand complexes (top three identified ligands U19, U94, U97 and reference ligand 89m) along with APO.

#### 3.3.2 Residue flexibility analysis

To understand the overall residue-wise protein dynamics in the systems, the evaluation of root mean square fluctuations (RMSF) plays a very important role. Therefore, to check the stability and flexibility of the residues, we have calculated the RMSFs for an entire simulation run of 200 ns using the *gmx_rmsf* module of GROMACS (Figure 4D-4F). From the RMSF profile, it can be understood that 1PK2 protein docked with ligands P27, P67, P76 exhibited a similar pattern of fluctuation like the reference ligand ACA at active site residues TRP80, THR79, LEU78, ARG 71, TRP69, ASP63 respectively. 1PK2 docked with ligands P27, P67 and P76 show a fluctuation peak of values 0.15nm, 0.16nm, 0.18 nm at residue VAL40 respectively but a higher fluctuation peak of 0.36 nm is observed at residue VAL40 for the reference ligand. This indicates that the ligands P27, P67, P76 are contributing to greater rigidity to the binding pocket of 1PK2 (Figure 4D). Similarly, 4CIK protein docked with ligands C19, C90, C97 exhibited a similar pattern of fluctuation like the reference ligand XO3 at active site residues ARG35, TRP62, TYR64, ARG71, TYR72 respectively. Likewise, 4CIK docked with ligands C19, C90, C97 exhibited fluctuation of 0.15 nm at the residues ASP55, ASP57 but in the case of reference ligand XO3, a higher fluctuation of 0.3 nm is observed. Also, XO3 shows a higher peak of 0.42nm at residues ASN9 and LYS8. This suggests that the identified ligands C19, C90, C97 have made the binding pocket more rigid and intact (Figure 4E). In the same manner, 5UGG protein docked with ligands U19, U94, U97 displayed the fluctuation pattern resembling the reference ligand 89M at active site residues LYS67, PHE68, CYX83, HID87, GLN239 respectively. Likewise, 5UGG docked with ligands U19, U94, U97 exhibited fluctuation of 0.2nm, 0.2nm, 0.28nm at the residues GLY762, SER765, CYS763 respectively, but in the case of reference ligand 89M, a higher fluctuation of 0.32 nm is observed. Also, 89M shows a higher peak of 0.5nm at residues TRP736, THR737, and GLY738. This propounds that the identified ligands U19, U94, U97 have made the binding pocket more inflexible (Figure 4F). From the RMSF profile of all the systems, it can be inferred that the ligands P27, P67, P76, C19, C90, C97, U19, U94, U97 are responsible for the restricted dynamics at the active site residue pocket of their respective protein targets and for imparting structural rigidity.

#### 3.3.3 Compactness analysis

To study the compactness of all the protein-ligand complexes, the radius of gyration (Rg) is computed using the *gmx_gyrate* function of GROMACS for a simulation time of 200ns. Rg is explained as the distance measured between the terminal end of protein and its center of mass during simulation. When a ligand docks to a protein, there is a conformational shift that changes the radius of gyration (Seeliger & De Groot, 2010). The compact protein structure tends to maintain a low Rg average deviation thus showing dynamic stability. The combined Rg plot of all ligands is represented in Figure 5A-5C. As provided in Figure 5A and supplementary table ST8, 1PK2 protein docked with ligands ACA (reference ligand), P27, P67, P76 shows an average Rg deviation of 1.31± 0.02 nm, 1.29± 0.01 nm, 1.27± 0.01 nm, 1.28 ± 0.02 nm respectively from 30-200 ns simulation time. Similarly, 4CIK protein docked with ligands X03 (reference ligand), C19, C90, C97 shows an average Rg deviation of 1.26 ± 0.01 nm, 1.23± 0.01 nm, 1.22± 0.01 nm, 1.21 ± 0.01 nm respectively from 30-200 ns simulation time (Figure 5B and supplementary table ST8). Likewise, 5UGG protein docked with ligands 89M (reference ligand), U19, U94, U97 exhibited an average Rg deviation of 1.77± 0.01 nm, 1.74± 0.01 nm, 1.75± 0.01 nm, 1.73 ± 0.01 nm respectively from 30-200 ns simulation time (Figure 5C and supplementary table ST8). Collectively, from the radius of gyration profile, it can be concluded that the identified ligands P27, P67, P76, C19, C90, C97, U19, U94, U97 are responsible for making the structure more compact compared to the reference ligands ACA, XO3, and 89M respectively.

**Figure 5:**
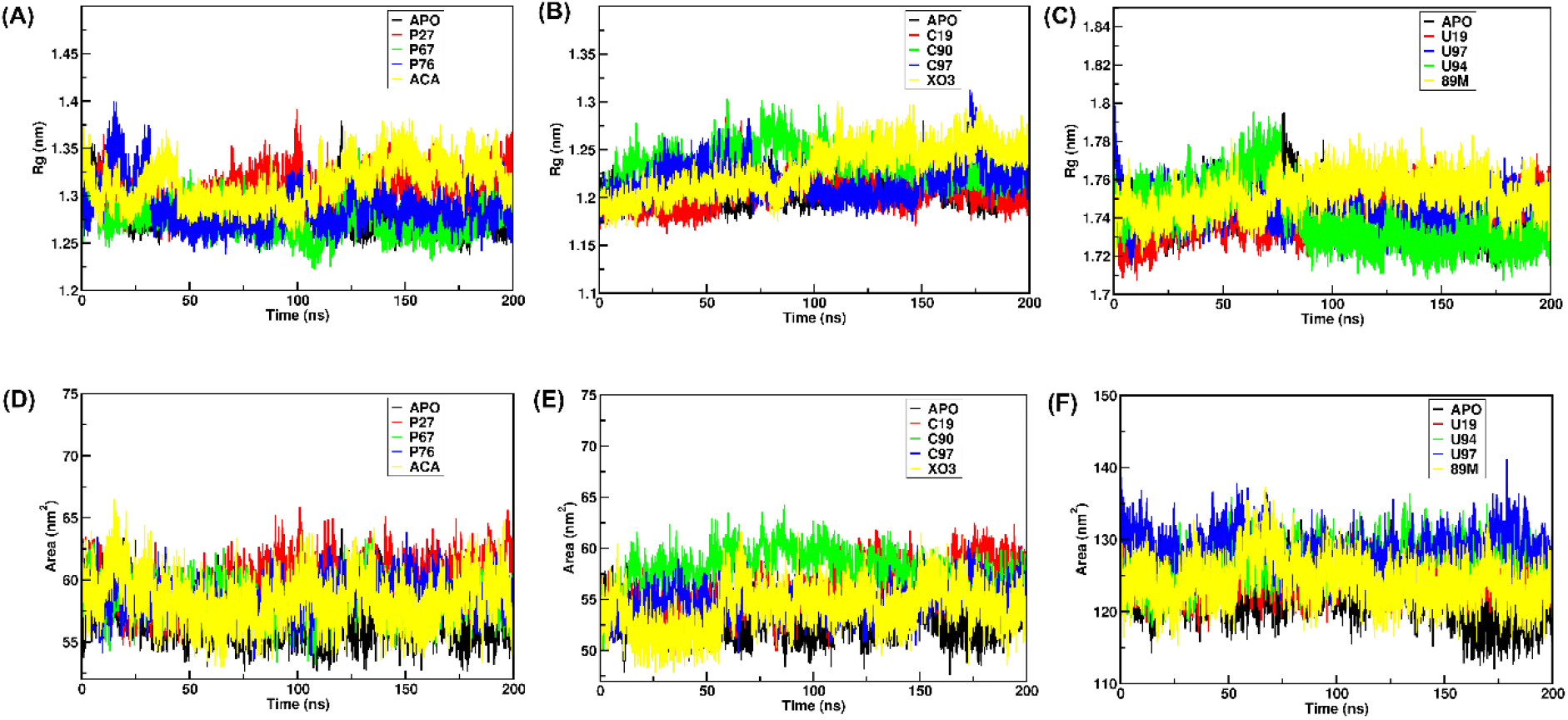
(A) Radius of gyration profile of 1PK2 protein-ligand complexes (top three identified ligands P27, P67, P76, and reference ligand ACA) along with APO. (B) The radius of gyration profile of 4CIK protein-ligand complexes (top three identified ligands C19, C90, C97and reference ligand XO3) along with APO. (C) The radius of gyration profile of 5UGG protein-ligand complexes (top three identified ligands U19, U94, U97, and reference ligand 89m) along with APO. (D) SASA profile of 1PK2 protein-ligand complexes (top three identified ligands P27, P67, P76, and reference ligand ACA) along with APO. (E) SASA profile of 4CIK protein-ligand complexes (top three identified ligands C19, C90, C97and reference ligand XO3) along with APO. (F) SASA profile of 5UGG protein-ligand complexes top three identified ligands U19, U94, U97 and reference ligand 89m) along with APO.

#### 3.3.4 Solvent Accessible Surface Area analysis

To calculate the solvent-accessible surface area (SASA) of a molecular surface, a method is employed which involves the *in-silico* rolling of a spherical probe that approximates a water molecule, encompassing a full-atom protein model (Durham et al., 2009). This procedure takes into account the expansion of the van der Waals radius for each atom by 1.4 Å (the radius of a polar solvent probe) followed by the calculation of the surface area of these atoms with expanded radius. Therefore, the SASA values of all the complexes were computed using the *gmx_sasa* function of GROMACS for a simulation time of 200 ns (Figure 5D-5F).

As provided in Figure 5D and supplementary table ST9, 1PK2 protein docked with ligands ACA (reference ligand), P27, P67, P76 shows an average SASA value of 58.94± 1.7 nm^2,^ 58.67± 1.83 nm^2^, 58.34± 1.54 nm^2^, 58.76± 1.53nm^2^ respectively from 30-200 ns simulation timeframe. It is observed that the SASA curves followed the same trend as the reference ligand ACA. Similarly, 4CIK protein docked with ligands X03 (reference ligand), C19, C90, C97 shows an average SASA value of 54.46 ± 2.0 nm^2^, 54.18± 2.3 nm^2^, 54.28± 1.72 nm^2^, 53.21± 1.7 nm^2^ respectively from 30-200 ns simulation time frame (Figure 5E and supplementary table ST9). The C19, C90, C97 show a similar pattern of average SASA values when compared to XO3. Likewise, 5UGG protein docked with ligands 89M (reference ligand), U19, U94, U97 exhibited an average SASA value of 124.75± 2.85 nm^2,^ 123.61± 2.36 nm^2^, 124.32± 2.62 nm^2^,124.24± 2.84 nm^2^ respectively from 30-200 ns simulation time frame (Figure 5F and supplementary table ST9). The same pattern of SASA values was obtained for all the identified ligands. It is observed that similar to reference ligands, the identified ligands make the active site residues of the protein targets readily accessible to the solvent surface, thus exposing the proteins to the hydration shell. Also, the SASA values obtained from the analysis of proteins docked with our identified ligands are found to be significantly similar to the reference protein-ligand complexes.

#### 3.3.5 Hydrogen Bond and Bond distribution analysis

The hydrogen bonds (H-bonds) are known to play a key role in forming stable contacts between the ligands and protein targets (Chen et al., 2016). In this study, H-bond analysis was performed on all the protein-ligand systems for a total simulation run of 200ns. The number of H-bonds and their bond frequencies are plotted using the *<u>gmx_hbon</u>d* tool of GROMACS and are represented in Figure 6A-6F. It is observed that the ligand P67 formed one stable H-bond of length 0.27 nm with the protein target 1PK2, and its frequency to form this bond is 20 % (Figure 6A). Likewise, the P76 formed 2-3 H-bonds during the simulation period and its frequency to form a bond of length 0.28 nm is 17.5% (Figure 6A). Similarly, P27 had formed a bond of length 0.29 nm with a frequency of 17% (Figure 6A). The reference ligand ACA did not form stable H-bonds and the frequency to form a bond of length 0.28 nm is only 10% which is very less compared to identified ligands (Figure 6A). In the case of the 4CIK protein target, the ligand C97 established stable 1-2 H-bonds during the simulation period and had a 23% frequency to form a bond of length 0.28nm (Figure 6B). In the same way, the C90 exhibited 1-2 H-bonds during the simulation period, and its ability to form a bond of length 0.27 is 18% (Figure 6B). Similarly, the ligand C19 formed 1 H-bond, and its frequency to form a bond of length 0.28 nm is 18% (Figure 6B). Lastly, we observed that the reference ligand XO3 formed unstable H-bonds during the simulation, and its frequency to form a bond of length 0.26 nm is only 10%, which is less compared to identified ligands (Figure 6B). Likewise, in the case of the 5UGG protein target, the ligand U97 formed 1-2 stable H-bonds during the simulation time, and its frequency to form a bond of length 0.27 nm is 22% (Figure 6C). The U94 ligand formed 2-4 stable H-bonds and the frequency to form a bond of length 0.27 nm is 27% (Figure 6C). Also, the ligand U19 formed 2-4 H-bonds, and its frequency to form a bond of length 0.28 nm is 20% (Figure 6C). Lastly, the reference ligand 89M formed 1-2 unstable H-bonds, and the frequency to form a bond of length nm is 15% (Figure 6C). Overall, from the H-bond analysis, it can be concluded that our identified ligands form stable and stronger H-bonds of length ≥ 0.28 nm with their target proteins compared to reference ligands.

**Figure 6:**
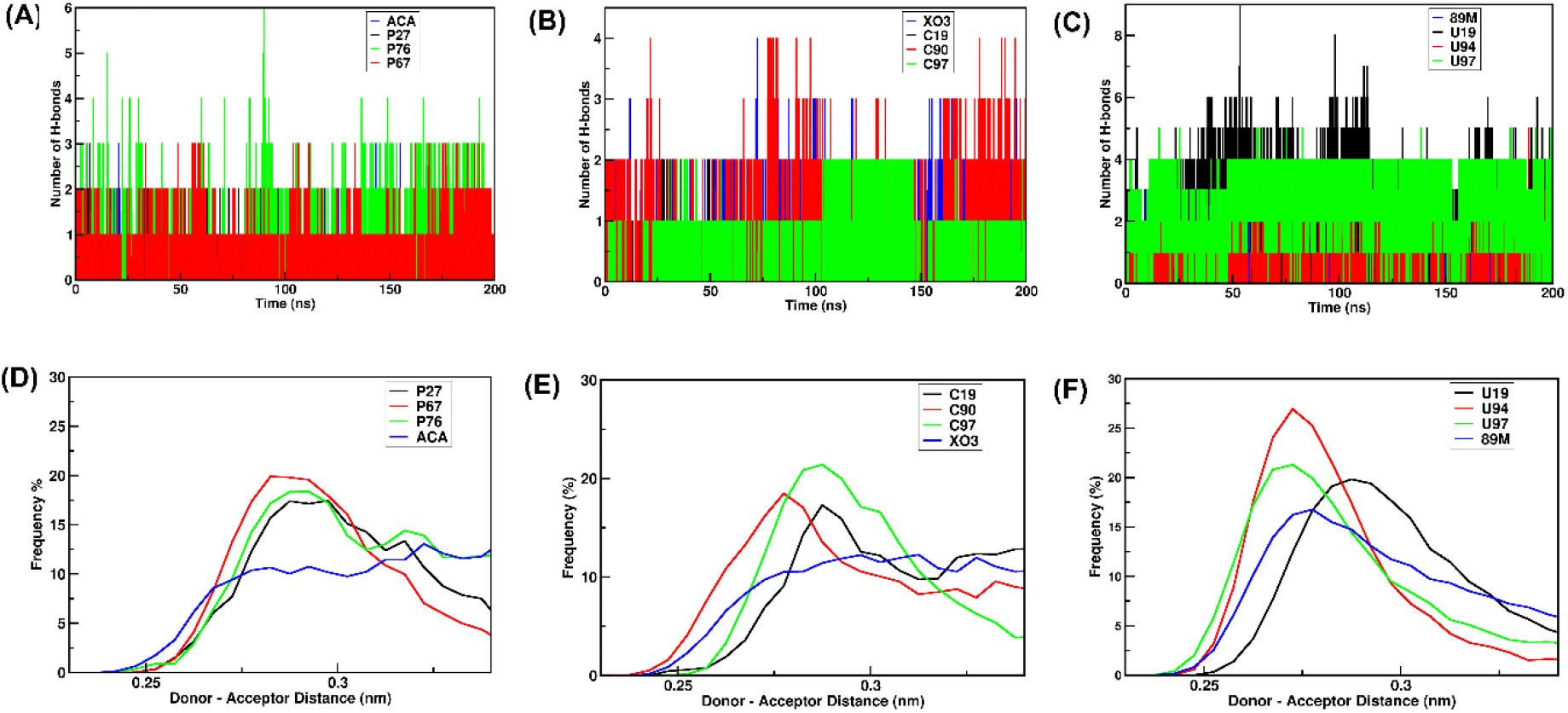
(A) Hydrogen bonding pattern of 1PK2 protein-ligand complexes (top three identified ligands P27, P67, P76, and reference ligand ACA) along with APO. (B) Hydrogen bonding pattern of 4CIK protein-ligand complexes (top three identified ligands C19, C90, C97and reference ligand XO3) along with APO. (C) Hydrogen bonding pattern of 5UGG protein-ligand complexes (top three identified ligands U19, U94, U97, and reference ligand 89m) along with APO. (D) H-Bond frequencies of 1PK2 protein-ligand complexes (top three identified ligands P27, P67, P76, and reference ligand ACA) along with APO. (E) H-Bond frequencies of 4CIK protein-ligand complexes (top three identified ligands C19, C90, C97 and reference ligand XO3) along with APO. (F) H-Bond frequencies of 5UGG protein-ligand complexes top three identified ligands U19, U94, U97 and reference ligand 89m) along with APO.

#### 3.3.6 Principal component analysis

In general, a protein-ligand complex that occupies a smaller phase space with a stable cluster denotes a highly stable complex (Papaleo et al., 2009). Therefore, to explore this phenomenon, we have performed Principal Component Analysis (PCA) on the trajectories obtained from the 200 ns simulation period for all the protein-ligand complexes as shown in the 2D plots of PCA (Figure 7A-7C). From the PCA graph of 1PK2, it is observed that the complexes formed by P27, P67, P76 with 1PK2 are clustered very closely and the area coverage is less compared to the reference ligand ACA (Figure 7A). In the same way, from the PCA graph of 4CIK, it is observed that the complexes formed by C19, C90, C97 with 4CIK are closely clustered and occupy less area compared to reference ligand XO3 (Figure 7B). Similarly, from the PCA graph of 5UGG, it is observed that the complexes formed by U19, U94, U97 with 5UGG are clustered and occupy small phase space compared to reference ligand 89M (Figure 7C). Thus, from the PCA analysis, it is implied that our identified ligands impart rigidity to their respective protein-ligand complexes by occupying smaller phase space with stable cluster formation compared to reference ligands.

**Figure 7:**
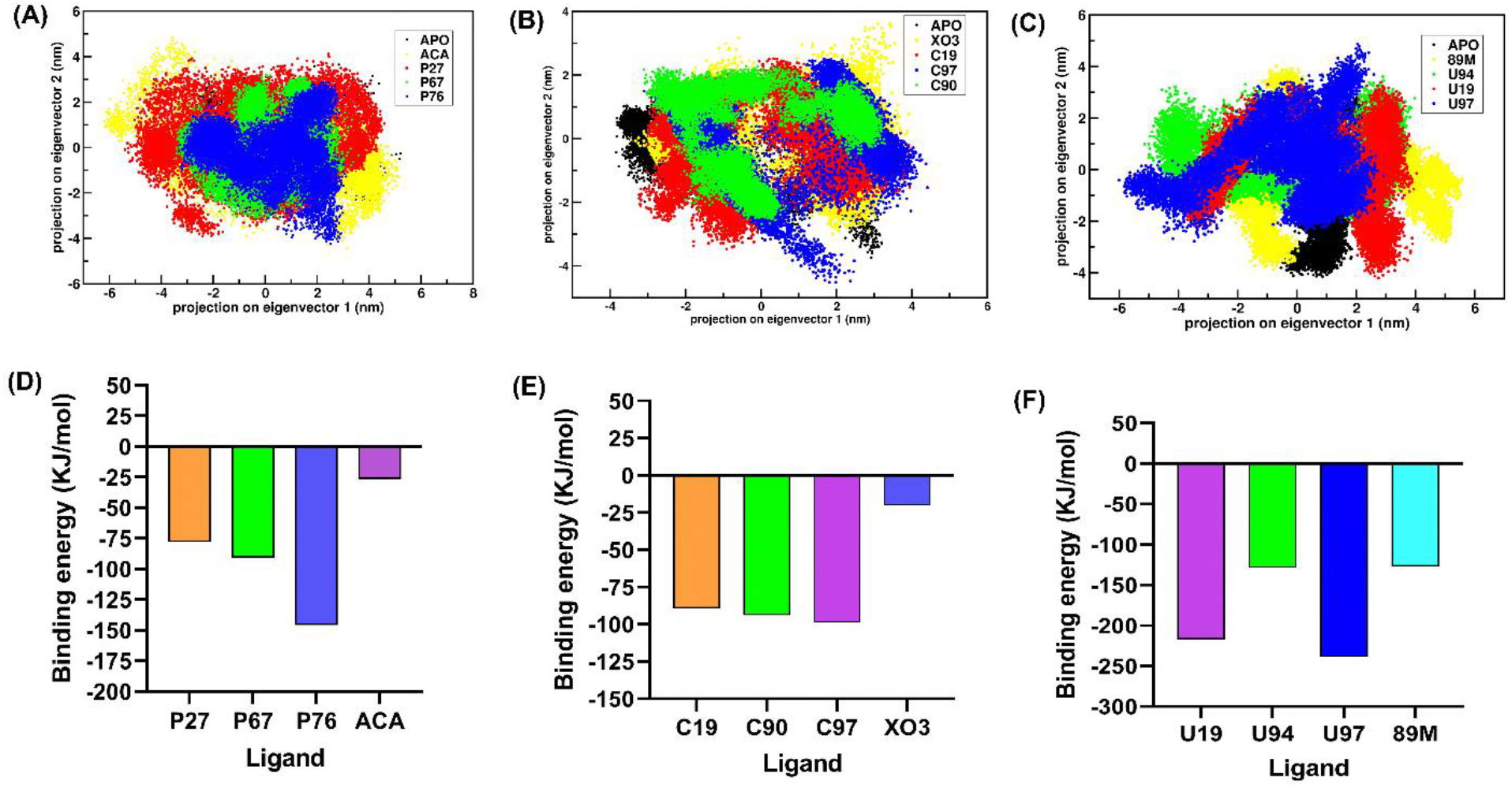
Principal component analysis and MMPBSA analysis. (A)Projection of motion of Apo-1PK2 in black; 1PK2-ACA complex in yellow; 1PK2-P27 complex in red; 1PK2-P67 in green; 1PK2-P76 in blue. (B) Projection of motion of Apo-4CIK in black; 4CIK-XO3 complex in yellow; 4CIK -C19 complex in red; 4CIK -C90 in green; 4CIK-C97 in blue. (C) Projection of motion of Apo-5UGG in black; 5UGG-89M complex in yellow; 5UGG -U19 complex in red; 5UGG -U94 in green; 5UGG -U97 in blue. (D) Binding energy profile (ΔG) of 1PK2 complexed with the identified ligands P27, P67, P76, and reference ligand ACA. (E) Binding energy profile (ΔG) of 4CIK complexed with the identified ligands C19, C90, C97, and reference ligand XO3. (F) Binding energy profile (ΔG) of 5UGG complexed with the identified ligands U19, U94, U97, and reference ligand 89M.

#### 3.3.7 Binding free energy calculation

The results obtained by calculation of the free energy of ligands give insights into ligands’ binding potential with bonded and non-bonded entities (Wang et al., 2018). In this study, the *g_mmpbsa* tool was used to calculate binding free energy (ΔG) for the last 50 ns of the simulation time (Figure 7D-7F). From the MMPBSA graphs, it is observed that the reference ligand ACA, and the identified ligands P27, P67, P76 bound to 1PK2 target protein exhibit binding energy of -26.460 kJ/mol and -77.975 kJ/mol, -90.798, -145.653 kJ/mol respectively (Figure 7D). Thus, the binding free energies (ΔG) of 1PK2 bound with our identified ligands were found to be significantly better than that of 1PK2 bound with the reference ligand. Similarly, from the MMPBSA graphs, it is observed that the reference ligand XO3 and the identified ligands C19, C90, C97 bound to the 4CIK protein target exhibit binding energy of - 20.066 kJ/mol, and -89.316 kJ/mol, -93.881 kJ/mol, -98.949 kJ/mol respectively (Figure 7E). Likewise, from the MMPBSA graphs, it is observed that the reference ligand 89M and the identified ligands U19, U94, U97 bound to the 5UGG protein target, exhibits binding energy of - 126.981 kJ/mol, and -217.150 kJ/mol, -168.674 kJ/mol, -238.846 kJ/mol respectively (Figure 7F). The 3D images of the top nine identified ligands P27, P67, P76, C19, C90, C97, U19, U94, U97 bound to the active site residues of their respective protein targets 1PK2, 4CIK, and 5UGG are shown in supplementary figure 1 [(A)-(I)]. Furthermore, from the binding energy calculation, it can also be concluded that out of the nine identified ligands, the top three ligands having the best binding energy are P76, C97, and U97 against their protein targets 1PK2, 4CIK, and 5UGG respectively. The 3D images of these top three ligands P76, C97, and U97 bound to the active site residues of their respective protein targets 1PK2, 4CIK, and 5UGG are shown in Figure 8 [(A)-(C)]. From the MMPBSA profile, it can be inferred that the identified ligands docked with their respective target protein have the highest electrostatic energy, SASA energy, and Vander Waal energy in comparison with reference ligands. This also indicates that despite intermittent hydrogen bond formations as observed during h-bond analysis, the ligands were conserved in the binding pocket by nonbonded interactions (electrostatic, polar, and nonpolar). Based on overall MD analysis and MMPBSA energies, we have found the ligands P76, C97. and U97 to be the best performing amongst the top nine ligands.

**Figure 8:**
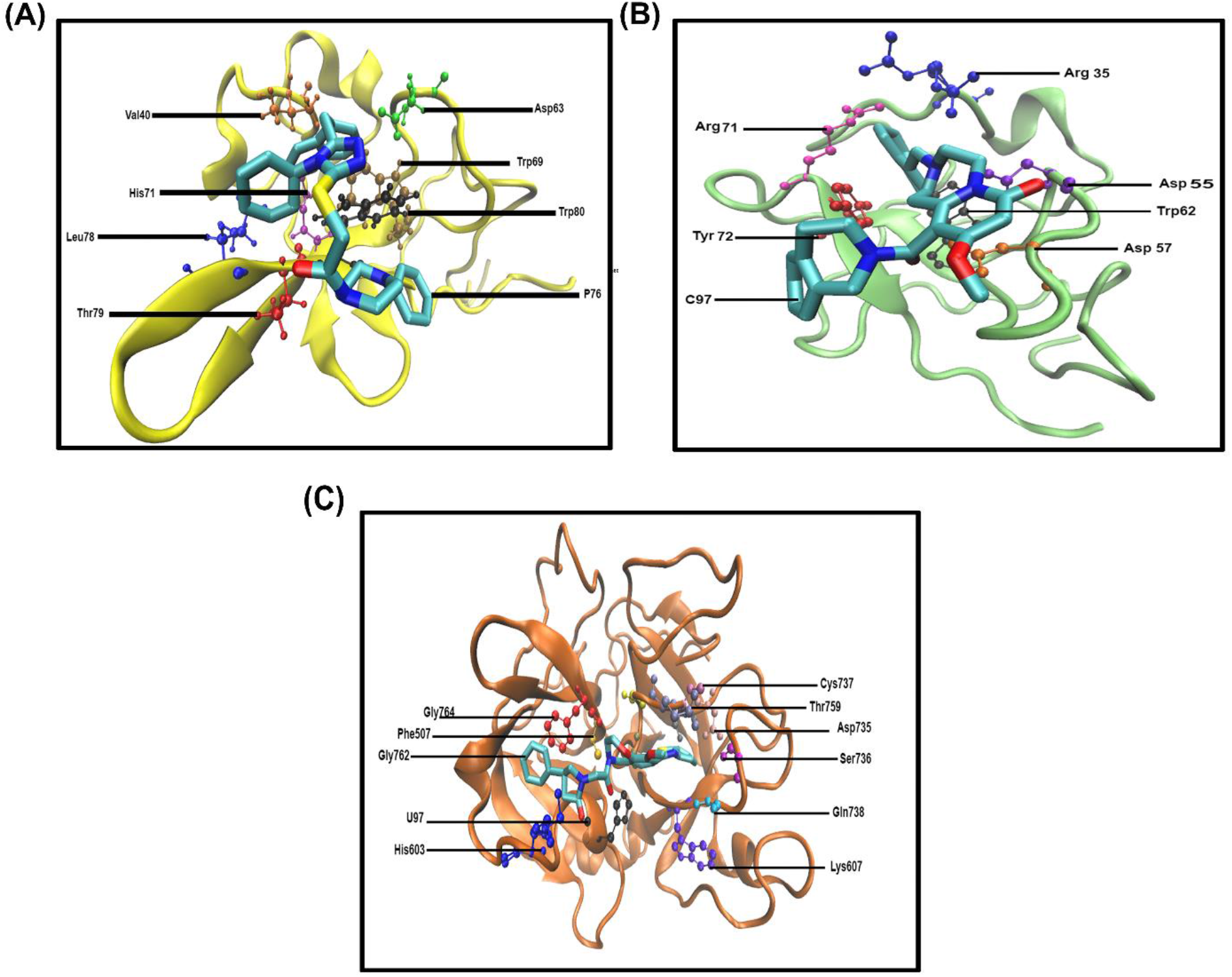
Schematic representation of the top three identified ligands docked to the active site residues of their respective protein targets using VMD. (A) P76 bound to 1PK2 (B) C97 bound to 4CIK (C) U97 bound to 5UGG. Proteins are shown in the New Cartoon representation. Active site residues of the proteins are shown in licorice representation. Identified ligands are shown in CPK representation.

## 4. Conclusions

In this study, we have identified compounds that can inhibit fibrinolysis by targeting each of the three important protein domains: the kringle-2 domain of tissue-type plasminogen activator, the kringle-1 domain of plasminogen, and the serine protease domain of plasminogen respectively. The top nine ligands (P27, P67, P76, C19, C90, C97, U19, U94, U97) were obtained by screening one million molecules from the ZINC database. These ligands were docked to their respective protein targets (1PK2, 4CIK, and 5UGG) using Autodock Vina, Schrödinger Glide, and ParDOCK/BAPPL+. Docking scores of these top nine ligands were evaluated and their ADMET profiling was also performed keeping the reported ligands (ACA, XO3, and 89M) as reference. All the top nine ligands, with the best scores and good ADMET results, were individually subjected to MD simulation for a period of 200 ns followed by detailed trajectory analysis. RMSD profile suggests that all the protein-ligand complexes have attained stable trajectories and are well equilibrated. From the RMSF analysis of all the systems, it can be inferred that the identified ligands are responsible for the restricted dynamics at the active site residue pocket of their respective protein targets, thereby imparting structural rigidity to the protein compared to the reference ligands. Rg data of all the systems reveals that in comparison to reference ligands, our identified ligands contribute greater compactness to the structure of their respective protein targets. From the SASA analysis, it is evident that similar to reference ligands, the identified ligands make the active site residues of the protein targets well exposed to the solvent, thereby making them readily accessible. From the h-bond analysis and bond distribution, it was observed that the identified ligands tend to form stronger H-bonds of length ≥ nm with their target proteins compared to reference ligands. PCA analysis helps us to understand that in comparison to reference ligands our identified ligands occupy smaller phase space and form stable clusters, thus conferring more stability to the protein-ligand complexes. The results obtained by analyzing molecular docking and MD simulation data are well-validated by the binding free energy calculation. From the MMPBSA profile, it is observed that, in the identified protein-ligand complexes, there exist strong interactions in terms of energy components other than hydrogen bonding such as electrostatic energy, SASA energy, and Vander Waal energy compared to reference protein-ligand complexes. Based on overall MD analysis and MMPBSA energies, we have found the ligands P76, C97, and U97 to be the best performing amongst the top nine ligands. Hence, our study strongly substantiates the point that our identified ligands can act as strong inhibitors and further experimental studies can be conducted to prove the therapeutic potential of these promising antifibrinolytic agents.

## Authorship Contribution

Suparna Banerjee: Investigation, Data curation, Formal analysis, Validation, Visualization, Writing - original draft, review& editing. Yeshwanth M.: Investigation, Data curation, Formal analysis, Validation, Visualization, Writing - original draft, review& editing. D. Prabhu: Data curation, Formal analysis, Visualization, review& editing. K. Sekar: Supervision, Prosenjit Sen: Conceptualization, Supervision, Writing - review& editing. Suparna Banerjee and Yeshwanth M. have contributed equally to the work.

## Acknowledgments

The authors owe sincere thanks to the Department of Science and Technology, Government of India, and Indian Association for the Cultivation of Science, Kolkata, India for providing all the necessary facilities. The authors acknowledge the HPC facilities offered by the centre of excellence in structural biology and bio-computing funded by the Department of Biotechnology, Government of India, and the Department of Computational and Data Sciences, Indian Institute of Science, Bangalore, India. Suparna Banerjee gratefully acknowledges the support of the Council of Scientific & Industrial Research, Govt. of India for fellowship.

