## Supplementary file for "Identification of potent anti-fibrinolytic compounds against plasminogen and tissue-type plasminogen activator using computational approaches"

**Running Title:** Identification of potent antifibrinolytic compounds against plasminogen and tissue-type plasminogen activator

Suparna Banerjee*[1], Yeshwanth. M*[2], Dhamodharan Prabhu[3], Kanagaraj Sekar[2] and Prosenjit Sen[#1]

[1]School of Biological Sciences, Indian Association for the Cultivation of Science, Jadavpur, Kolkata 700032, West Bengal, India

[2] Department of Computational and Data Sciences, Indian Institute of Science, Bangalore 560012, Karnataka, India

[3]Research and Development Wing, Sree Balaji Medical College and Hospital-BIHER, Chennai 600044, Tamil Nadu, India

* These authors have made equal contribution, and should be treated as co-first authors

Supplementary Table ST1: AUTODOCK-VINA scores of top 10 identified ligands docked with their respective protein targets.

| **AUTODOCK-VINA SCORES (TOP 10)** | | | | | |
|---|---|---|---|---|---|
| 1PK2 | | 4CIK | | 5UGG | |
| Ligands | Binding energy (kcal/mol) | Ligands | Binding energy (kcal/mol) | Ligands | Binding energy (kcal/mol) |
| **P76** | **-8.9** | **C97** | **-8.8** | **U97** | **-11.4** |
| **P27** | **-8.3** | C38 | -8.6 | **U94** | **-11** |
| P95 | -7.8 | **C19** | **-8.5** | **U19** | **-10.9** |
| **P67** | **-7.6** | **C90** | **-8.2** | U90 | -10.63 |
| P03 | -7.58 | C63 | -8.1 | U26 | -10.6 |
| P83 | -7.57 | C20 | -8 | U08 | -10.5 |
| P09 | -7.54 | C07 | -7.9 | U51 | -10.3 |
| P01 | -7.51 | C10 | -7.8 | U80 | -10.2 |
| P71 | -7.4 | C27 | -7.7 | U18 | -10 |
| P78 | -7.32 | C66 | -7.6 | U87 | -9.88 |
| ACA-std | -3.9 | XO3-std | -7.4 | 89M-std | -9 |

Supplementary Table ST2: PARDOCK/BAPPL+ scores of top 10 identified ligands docked with their respective protein targets.

| **PARDOCK/BAPPL+ SCORES (TOP 10)** | | | | | |
|---|---|---|---|---|---|
| 1PK2 | | 4CIK | | 5UGG | |
| Ligands | Binding energy (kcal/mol) | Ligands | Binding energy (kcal/mol) | Ligands | Binding energy (kcal/mol) |
| P66 | -11.85 | C86 | -11.03 | **U97** | **-8.33** |
| P74 | -11.09 | C62 | -10.76 | U7 | -8.08 |
| P79 | -10.41 | C83 | -10.71 | **U94** | **-8.02** |
| P78 | -10.18 | C22 | -10.68 | U59 | -8.01 |
| P44 | -9.98 | C38 | -10.6 | **U19** | **-7.9** |
| P48 | -9.85 | C12 | -10.57 | U55 | -7.86 |
| P22 | -9.73 | **C97** | **-9.47** | U15 | -7.74 |
| **P76** | **-9.63** | C18 | -9.2 | U81 | -7.7 |
| **P27** | **-9.6** | **C19** | **-9** | U35 | -7.69 |
| **P67** | **-8.84** | **C90** | **-8.9** | U40 | -7.6 |
| ACA-std | -4.9 | XO3-std | -6.66 | 89M-std | -5.9 |

Supplementary Table ST3: GLIDE scores of top 10 identified ligands docked with their respective protein targets.

| GLIDE SCORES (TOP 10) | | | | | |
|---|---|---|---|---|---|
| 1PK2 | | 4CIK | | 5UGG | |
| Ligands | Binding energy (kcal/mol) | Ligands | Binding energy (kcal/mol) | Ligands | Binding energy (kcal/mol) |
| P82 | -10.11 | C32 | -9.9 | U88 | -9.78 |
| P86 | -9.98 | C62 | -9.5 | **U97** | **-9.6** |
| P67 | -9.7 | **C97** | **-9.3** | U96 | -9.53 |
| P48 | -9.56 | **C19** | **-8.92** | **U94** | **-9.41** |
| **P76** | **-9.42** | C74 | -8.81 | U49 | -9.33 |
| **P27** | **-9.31** | **C90** | **-8.75** | U22 | -9.21 |
| **P67** | **-8.6** | C36 | -8.56 | **U19** | **-8.87** |
| P22 | -8.4 | C49 | -8.51 | U11 | -8.76 |
| P64 | -8.37 | C40 | -8.42 | U09 | -8.54 |
| P93 | -8.1 | C15 | -8.13 | U08 | -8.39 |
| ACA-std | -7.25 | XO3-std | -7.4 | 89M-std | -7.5 |

Supplementary Table ST4: GLIDE scores of top 3 identified ligands docked with their respective protein targets.

| GLIDE DOCKING SCORES (TOP 3 identified and reference ligands) | | | | | |
|---|---|---|---|---|---|
| Protein target | Ligand | Docking Score | XP Gscore | Glide Gscore | Glide energy |
| 1PK2 | P27 | -9.12 | -9.42 | -9.42 | -39.63 |
| | P67 | -8.52 | -8.6 | -8.6 | -37.27 |
| | P76 | -9.31 | -9.31 | -9.31 | -35.56 |
| | ACA | -7.11 | -7.25 | -7.25 | -22.85 |
| 4CIK | C19 | -8.34 | -8.92 | -8.92 | -43.23 |
| | C90 | -7.82 | -8.75 | -8.75 | -39.23 |
| | C97 | -8.9 | -9.3 | -9.3 | -40.01 |
| | XO3 | -6.75 | -7.15 | -7.15 | -31.62 |
| 5UGG | U19 | -9.21 | -9.6 | -9.6 | -88.73 |
| | U94 | -8.67 | -9.41 | -9.41 | -85.92 |
| | U97 | -8.33 | -8.87 | -8.87 | -79.37 |
| | 89M | -7.45 | -7.5 | -7.5 | -70.75 |

Supplementary Table ST5: ADMET properties of top 3 identified ligands for each protein target.

| Protein | Ligand name | Solubility level | BBB Penetration level | Absorption level | Extension CYP2D6 | Extension hepatotoxicity |
|---|---|---|---|---|---|---|
| | P27 | 1 | 3 | 0 | -1.52 (false) | -11.08 (false) |
| | P67 | 2 | 3 | 1 | -9.95 (false) | -18.01 (false) |
| | P76 | 2 | 3 | 1 | -7.24 (false) | -6.26 (false) |
| 1PK2 | ACA | 5 | 3 | 0 | -4.03 (false) | -11.83 (false) |
| | C19 | 2 | 3 | 1 | -0.13 (false) | -14.09 (false) |
| | C90 | 2 | 3 | 0 | -4.45 (false) | -7.81 (false) |
| | C97 | 3 | 3 | 0 | -1.8 (false) | -4.85 (false) |
| 4CIK | XO3 | 4 | 3 | 0 | -4.11 (false) | -6.21 (false) |
| | U19 | 2 | 4 | 2 | -3.21 (false) | -7.68 (false) |
| | U94 | 1 | 3 | 1 | -7.24 (false) | -5.11 (false) |
| | U97 | 2 | 4 | 0 | -7.28 (false) | -6.51 (false) |
| 5UGG | 89M | 2 | 4 | 0 | -9.51 (false) | -5.81 (false) |

ADMET properties: Extension hepatotoxicity: <1 is nontoxic. CYP2D6: -ve is noninhibitors and +ve is inhibition, human intestinal absorption level: 0 (good); 1 (moderate); 2 (low); 3 (very low), (aqueous solubility): 0 (extremely low); 1 (low); 2 (good); 3 (optimal); 4 (too soluble), BBB (blood brain barrier): 0(very high); 1(high); 2 (mediums); 3 (low); 4 (undefined).

Supplementary Table ST6: TOPKAT properties of top 3 identified ligands for each protein target.

| PROTEIN | Ligand name | NTP carcinogenicity call (male rat) (v3.2) | NTP carcinogenicity call (female rat) (v3.2) | Developmental toxicity potential (DTP) (v3.1) | Skin irritation (v6.1) | Ames mutagenicity (v3.1) |
|---|---|---|---|---|---|---|
| | P27 | NC | NC | NT | NI | NM |
| | P67 | NC | NC | NT | NI | NM |
| | P76 | NC | NC | NT | NI | NM |
| 1PK2 | ACA | NC | NC | T | NI | NM |
| | C19 | NC | NC | NT | NI | NM |
| | C90 | NC | NC | NT | NI | NM |
| | C97 | NC | NC | NT | NI | NM |
| 4CIK | XO3 | C | C | T | NI | NM |
| | U19 | NC | NC | NT | NI | NM |
| | U94 | NC | NC | NT | NI | NM |
| 5UGG | U97 | NC | NC | NT | NI | NM |

| | 89M | C | C | NT | I | M |
|---|---|---|---|---|---|---|

TOPKAT properties: NC -Non-carcinogenic, C-Carcinogenic, NT-Non-Toxic, T-Toxic, NI-Non-Irritant, I-Irritant, NM-Non-Mutagenic, M-Mutagenic

Supplementary Table ST7: RMSD values of all the protein-ligand complexes

| RMSD | | |
|---|---|---|
| Protein-Ligand Complex (30-200ns) | Average(nm) | SD -/+ |
| 1PK2-APO | 0.44 | 0.05 |
| 1PK2-D27 | 0.45 | 0.05 |
| 1PK2-D67 | 0.36 | 0.04 |
| 1PK2-D76 | 0.4 | 0.03 |
| 1PK2-ACA | 0.32 | 0.05 |
| 4CIK-APO | 0.34 | 0.06 |
| 4CIK-D19 | 0.29 | 0.03 |
| 4CIK-D90 | 0.3 | 0.03 |
| 4CIK-D97 | 0.34 | 0.04 |
| 4CIK-XO3 | 0.31 | 0.03 |
| 5UGG-APO | 0.27 | 0.02 |
| 5UGG-D19 | 0.29 | 0.02 |
| 5UGG-D94 | 0.3 | 0.02 |
| 5UGG-D97 | 0.31 | 0.02 |
| 5UGG-89M | 0.24 | 0.02 |

Supplementary Table ST8: Rg values of all the protein-ligand complexes

| Radius of gyration | | |
|---|---|---|
| Protein-Ligand Complex (30-200ns) | Average(nm) | SD -/+ |
| 1PK2-APO | 1.26 | 0.02 |
| 1PK2-D27 | 1.29 | 0.01 |
| 1PK2-D67 | 1.27 | 0.01 |
| 1PK2-D76 | 1.28 | 0.02 |
| 1PK2-ACA | 1.31 | 0.02 |
| 4CIK-APO | 1.2 | 0.01 |
| 4CIK-D19 | 1.23 | 0.01 |
| 4CIK-D90 | 1.22 | 0.01 |
| 4CIK-D97 | 1.21 | 0.01 |
| 4CIK-XO3 | 1.26 | 0.01 |
| 5UGG-APO | 1.74 | 0.01 |
| 5UGG-D19 | 1.74 | 0.01 |
| 5UGG-D94 | 1.75 | 0.01 |
| 5UGG-D97 | 1.73 | 0.01 |
| 5UGG-89M | 1.77 | 0.01 |

Supplementary Table ST9: SASA values of all the protein-ligand complexes

| SASA | | |
|---|---|---|
| Protein-Ligand Complex (30-200ns) | Average(nm ) | SD -/+ |
| 1PK2-APO | 56.9 | 1.86 |
| 1PK2-D27 | 58.67 | 1.83 |
| 1PK2-D67 | 58.34 | 1.54 |
| 1PK2-D76 | 58.76 | 1.53 |
| 1PK2-ACA | 58.94 | 1.7 |
| 4CIK-APO | 53.01 | 1.56 |
| 4CIK-D19 | 54.18 | 2.3 |
| 4CIK-D90 | 54.28 | 1.72 |
| 4CIK-D97 | 53.21 | 1.7 |
| 4CIK-XO3 | 54.46 | 2 |
| 5UGG-APO | 122.63 | 3.2 |
| 5UGG-D19 | 123.61 | 2.36 |
| 5UGG-D94 | 124.32 | 2.62 |
| 5UGG-D97 | 124.24 | 2.84 |
| 5UGG-89M | 124.75 | 2.85 |

Supplementary Figure 1: Schematic representation of the identified ligands P27and P67 docked to the active site residues of 1PK2 protein target using VMD. (A) P27 bound to 1PK2 (B) P67 bound to 1PK2. Proteins are shown in New Cartoon representation. Active site residues of the proteins are shown in licorice representation. Identified ligands are shown in CPK representation.

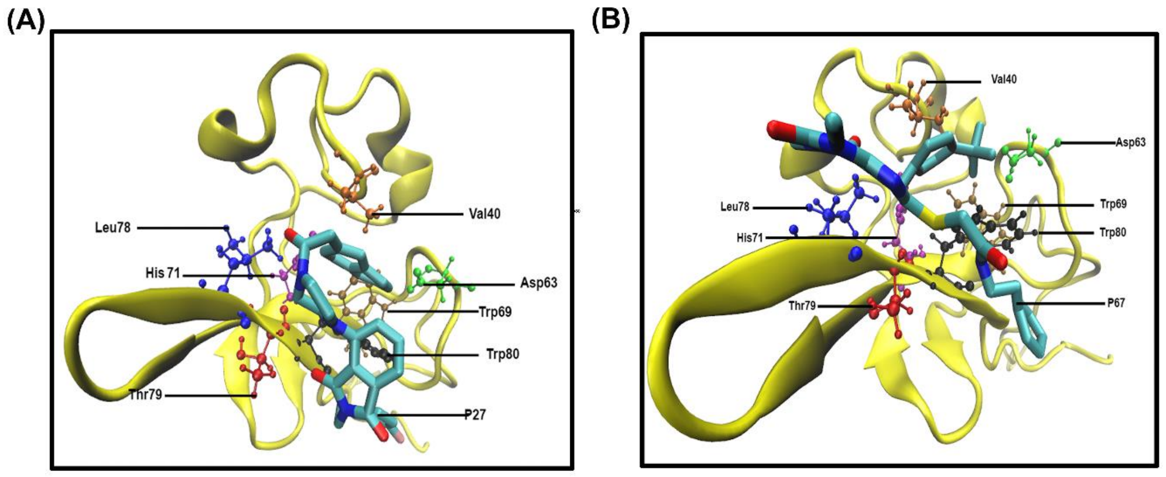

Supplementary Figure 2: Schematic representation of the identified ligands C19 and C90 docked to the active site residues of 4CIK protein target using VMD. (A) C19 bound to 4CIK (B) C90 bound to 4CIK. Proteins are shown in New Cartoon representation. Active site residues of the proteins are shown in licorice representation. Identified ligands are shown in CPK representation.

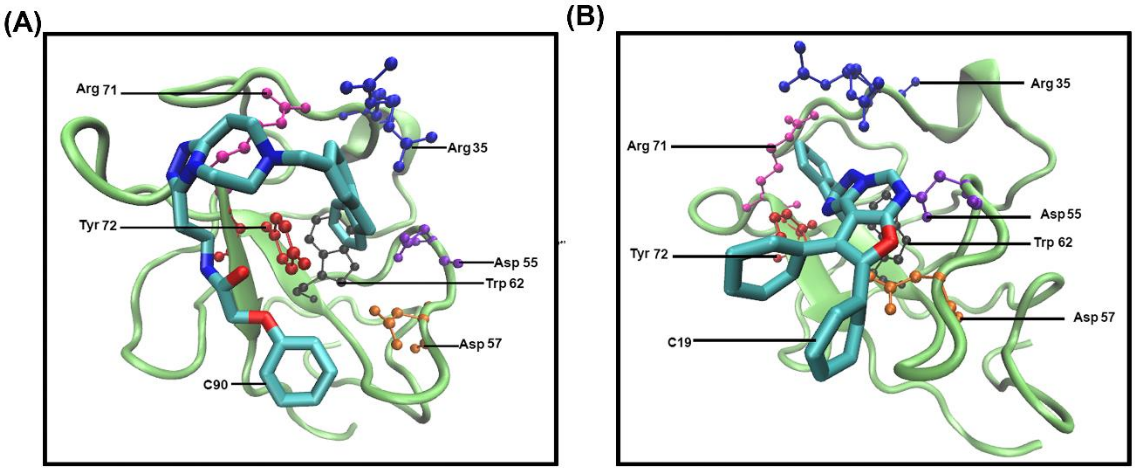


Supplementary Figure 3: Schematic representation of the identified ligands U90 and U19 docked to the active site residues of 5UGG protein target using VMD. (A) U90 bound to 5UGG (B) U97 bound to 5UGG. Proteins are shown in New Cartoon representation. Active site residues of the proteins are shown in licorice representation. Identified ligands are shown in CPK representation.

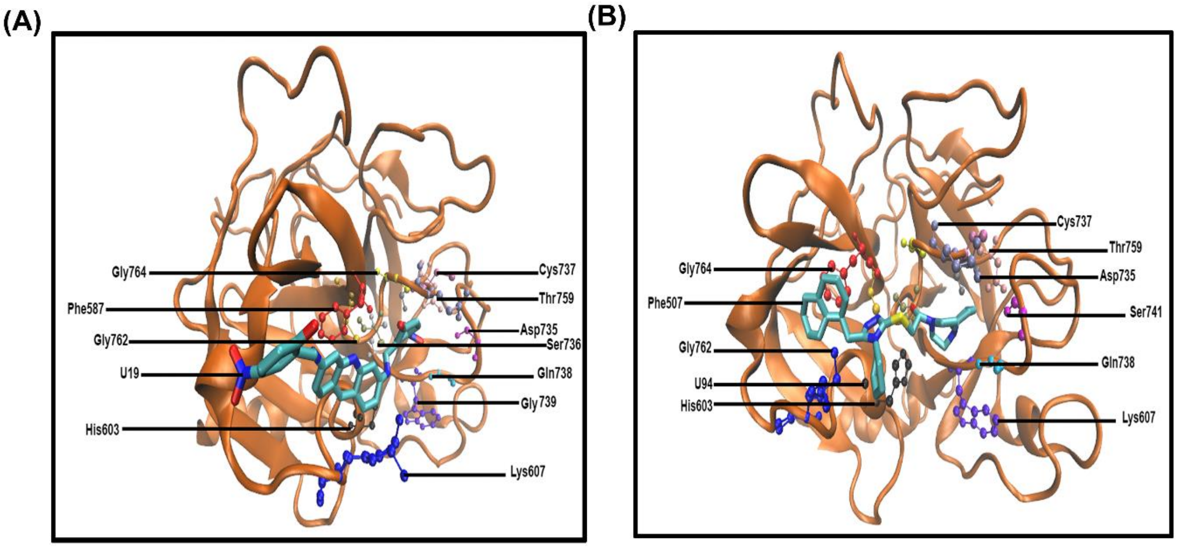
